# Silk-Ovarioids: Establishment and characterization of human ovarian primary cells 3D-model system

**DOI:** 10.1101/2024.07.31.606024

**Authors:** Valentina Di Nisio, Tianyi Li, Zhijie Xiao, Kiriaki Papaikonomou, Anastasios Damdimopoulos, Ákos Végvári, Filipa Lebre, Ernesto Alfaro-Moreno, Mikael Pedersen, Terje Svingen, Roman Zubarev, Ganesh Acharya, Pauliina Damdimopoulou, Andres Salumets

## Abstract

*In vitro* models that mimic ovaries are crucial for elucidating the biological mechanisms underlying follicle activation and growth. Three-dimensional (3D) systems are particularly relevant because they can replicate the heterogeneity and cell-cell communication between different ovarian cell types. However, complex models using human ovarian primary cells have not yet been established. In this study, we developed and characterized long-term cultured 3D models of primary ovarian somatic cells isolated from adult tissues, using Biosilk as a scaffold. We successfully established both ovarian cortex- and medulla-derived 3D systems, termed Silk-Ovarioids. The presence of key ovarian somatic cell types – including granulosa, stromal, endothelial, and perivascular cells – was confirmed by transcriptomics, proteomics, and immunostaining. Notably, Silk-Ovarioids exhibited the formation of a pro-angiogenic hypoxic core, as evidenced by the development of vessel-like structures after six weeks of culture. The Silk-Ovarioids demonstrated low intra-batch variability and long-term culture stability, underscoring their potential as a robust step towards creating a bioengineered, patient-specific artificial ovary.

## Introduction

A woman’s fertility potential relies on her ovarian reserve, also referred to as the follicular pool (La Marca *et al*, 2012). The ovarian reserve is defined as the number of dormant follicles in the ovarian cortex that can either be stimulated to grow and mature or undergo atresia. The ovarian reserve can be reduced by natural physiological factors (*e.g.*, age), pathological processes (*e.g.*, premature ovarian insufficiency, POI), and exposure to toxic chemicals or pharmaceuticals (*e.g.*, iatrogenic POI) (Esencan *et al*, 2022; Vabre *et al*, 2017; Bai & Wang, 2022; Mehedintu *et al*, 2021). Improvements in the treatment of childhood and young adult cancers and hematological diseases have led to increased survival rates. Consequently, the number of survivors living with long-term side effects like infertility and POI is rising (Donnez *et al*, 2006).

From a clinical perspective, ovarian tissue cryopreservation for future auto-transplantation is the only procedure available for prepubertal girls who are at very high risk of infertility from cancer treatment (Donnez *et al*, 2006; Antonouli *et al*, 2023). However, for malignancies like leukemias that are systemic, the high risk of reintroducing malignant cells constitutes a major concern and restricts the applicability of this approach (Diáz-Garciá *et al*, 2019). These cases necessitate the exploration of other resources, such as *in vitro* follicle growth and the collection of immature oocytes for further *in vitro* maturation in assisted reproductive technologies (ART) laboratories.

Obtaining *in vitro* matured metaphase II oocytes from *in vitro* grown follicles remains a challenge for both basic research and clinical implementation in ART programs (Hu *et al*, 2023; Barbato *et al*, 2023; Hao *et al*, 2020). In fact, *in vitro* follicle growth methodology has been developed mainly using animal models, and its application is still limited due to many drawbacks. These include the inability to maintain follicle architecture for long-term culture and the lack of standardization for translation to human application (Simon *et al*, 2020). Despite differences across multiple protocols for three-dimensional (3D) ovarian models, one commonality is the need to introduce or induce extracellular matrix (ECM) formation in culture (Hovatta *et al*, 1997; Shikanov *et al*, 2011; Desai *et al*, 2012). ECM is instrumental to the survival, growth, and establishment of a well-defined structure of somatic cells that are pivotal for mammalian *in vitro* follicle support (Grosbois *et al*, 2023). As reported in previous studies, the human ovarian tissue architecture is based on the expression of laminin, collagen, and other fibronectin types to support the different ovarian functions during the female reproductive lifespan (Hao *et al*, 2020; Ouni *et al*, 2021, 2022). Despite the advances obtained in the *in vitro* ovarian cortex tissue culture (Hao *et al*, 2024), this system displays a number of limitations that makes this golden standard insufficient for the next steps of *in vitro* folliculogenesis and ovarian somatic cells studies. Therefore, we are in need to explore new and diverse 3D models to overpass this bottleneck and implement ovarian culture systems for both somatic and germ cells investigation.

Aside from folliculogenesis-related research, new technologies have been explored for *in vitro* ovarian modeling to study physiopathological conditions, such as ovarian cancer. These technologies include the development of organ-on-a-chip systems using microfluidic devices (Yan *et al*, 2023) and the reconstruction of ovaries from pluripotent stem cells in mice (Yoshino *et al*, 2021). In addition, human-specific ovarian organoids represent a promising alternative to animal models with the added benefit of exploring personalized treatments using patient-specific cancer models (Nanki *et al*, 2020; Graham *et al*, 2023). Nevertheless, we still lack *in vitro* models of ovaries derived from healthy tissue that can be maintained in long-term culture and used to advance our understanding of ovarian biology and functions. A major reason for this is the limited access to human ovarian tissues needed for model development, as well as the difficulties in handling and growing ovarian cells.

To address these challenges, we aimed to develop robust 3D culture models that faithfully mimic the human ovarian somatic cell niche. We tested three different approaches using ovarian material from the same patients: matrix-free ovarian spheroids (MFOS), a Matrigel-based three-layer gradient system (3LGS), and Biosilk-based floating spheroids (Silk-Ovarioids) for the 3D culture of human ovarian primary somatic cells. Somatic cells from adult patients grew robustly only in the Biosilk system, where they formed Silk-Ovarioids that could be maintained in a floating culture for 6 weeks. Silk-Ovarioids harbored all key ovarian cell types and exhibited pro-angiogenic activity.

## Results

### Study setup

In this study, we compared three different approaches to establish a 3D model based on adult ovarian, primary somatic cells that closely mimics the natural follicle growth environment in humans: i) MFOS, ii) a Matrigel-based 3LGS in hanging drops, and iii) Silk-Ovarioids. Ovarian tissue samples retrieved from five different patients undergoing gender-affirming surgery were dissociated and used for the 3D *in vitro* cultures (Fig. 1). First, the survival of cells in the 3D cultures was compared to the cells from the same patients in 2D monolayer cultures. Where stable structures were formed (*e.g.*, endothelial marker-positive tubules), the quality of the structures was assessed in terms of cell survival/death and morphological and molecular assays (Fig. 1).

**Figure 1.**
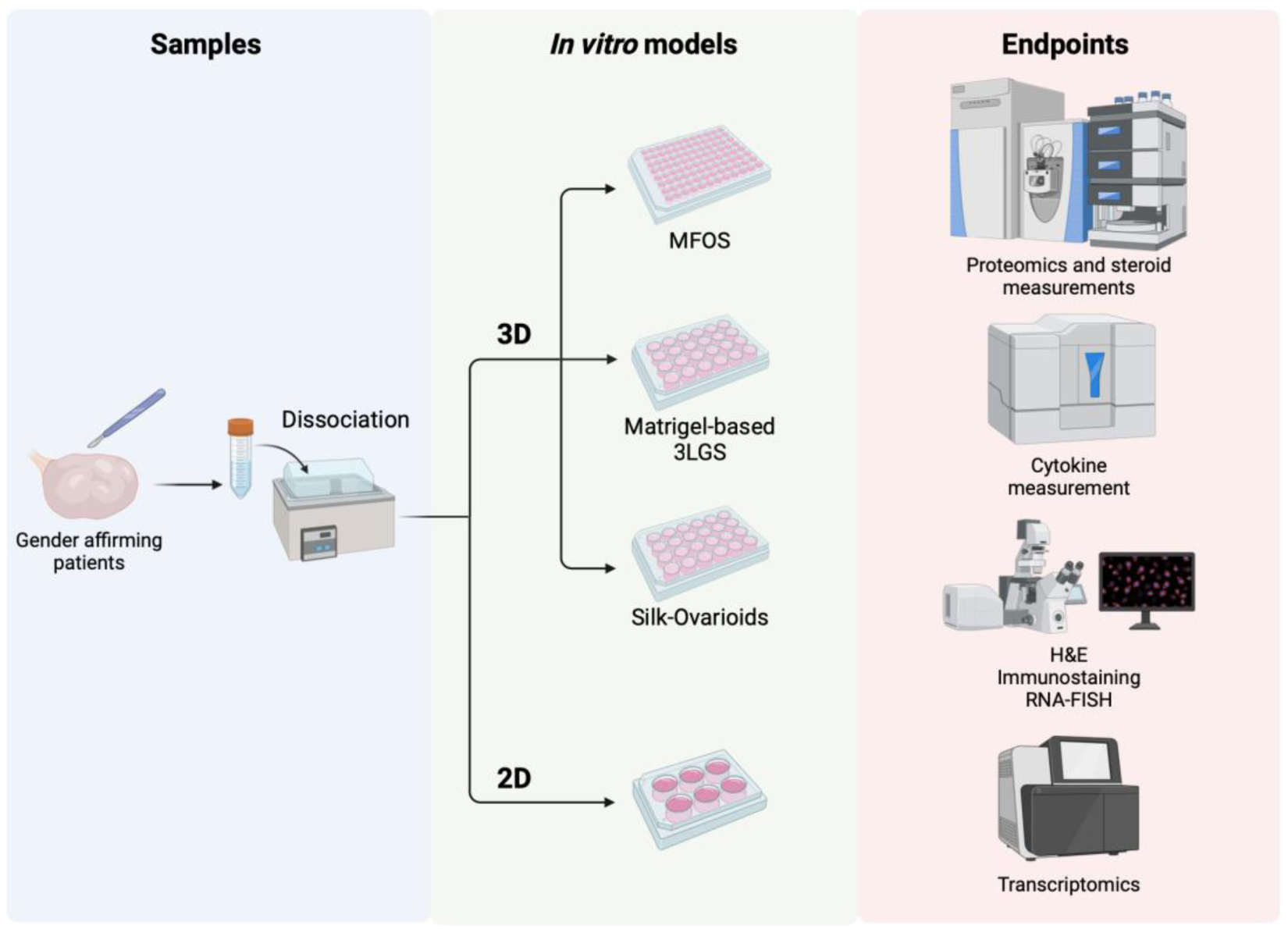
Overview of experimental setup. Experimental setup of 2D and 3D *in vitro* models of ovarian primary cells, and endpoints of the study. MFOS: matrix-free ovarian spheroids; 3LGS: three-layer gradient system; H&E: hematoxylin and eosin staining; RNA-FISH: RNA fluorescence *in situ* hybridization. Created with BioRender.com.

The most promising model, the Silk-Ovarioids, was further compared to the human ovarian cortex and medulla *via* transcriptomics and proteomics analyses, followed by validation of the bioinformatic analysis through immunostaining of the selected markers (Fig. 1). Finally, the function of the models was tested by analyzing the secretion of cytokines and steroids (Fig. 1).

### Approaches to grow adult ovarian somatic cells in 3D

Adult ovarian somatic cells from the cortex and medulla were cultured in three different models. We first assessed the feasibility of spheroid formation in matrix-free conditions by seeding cells at different densities (3 × 10^4^, 6 × 10^4^, 1.2 × 10^5^ cells/well).

MFOS of variable sizes (100-250 µm) could be obtained from low and medium cell densities without the support of any matrix in low attachment plates after eight days of culture; however, the structures spontaneously disintegrated after two weeks (Fig. 2a). Concurrently, we applied the Matrigel-based 3LGS culture, previously used to successfully generate rat and human testicular organoids (Oliver *et al*, 2021; Alves-Lopes *et al*, 2017, 2018), to ovarian primary cells. Small medulla-derived spheroids (100-200 µm) could be cultured for up to 11 days before the structures began to disaggregate, whereas, surprisingly, no spheroids formed using cells isolated from cortical tissue (Fig. 2b).

**Figure 2.**
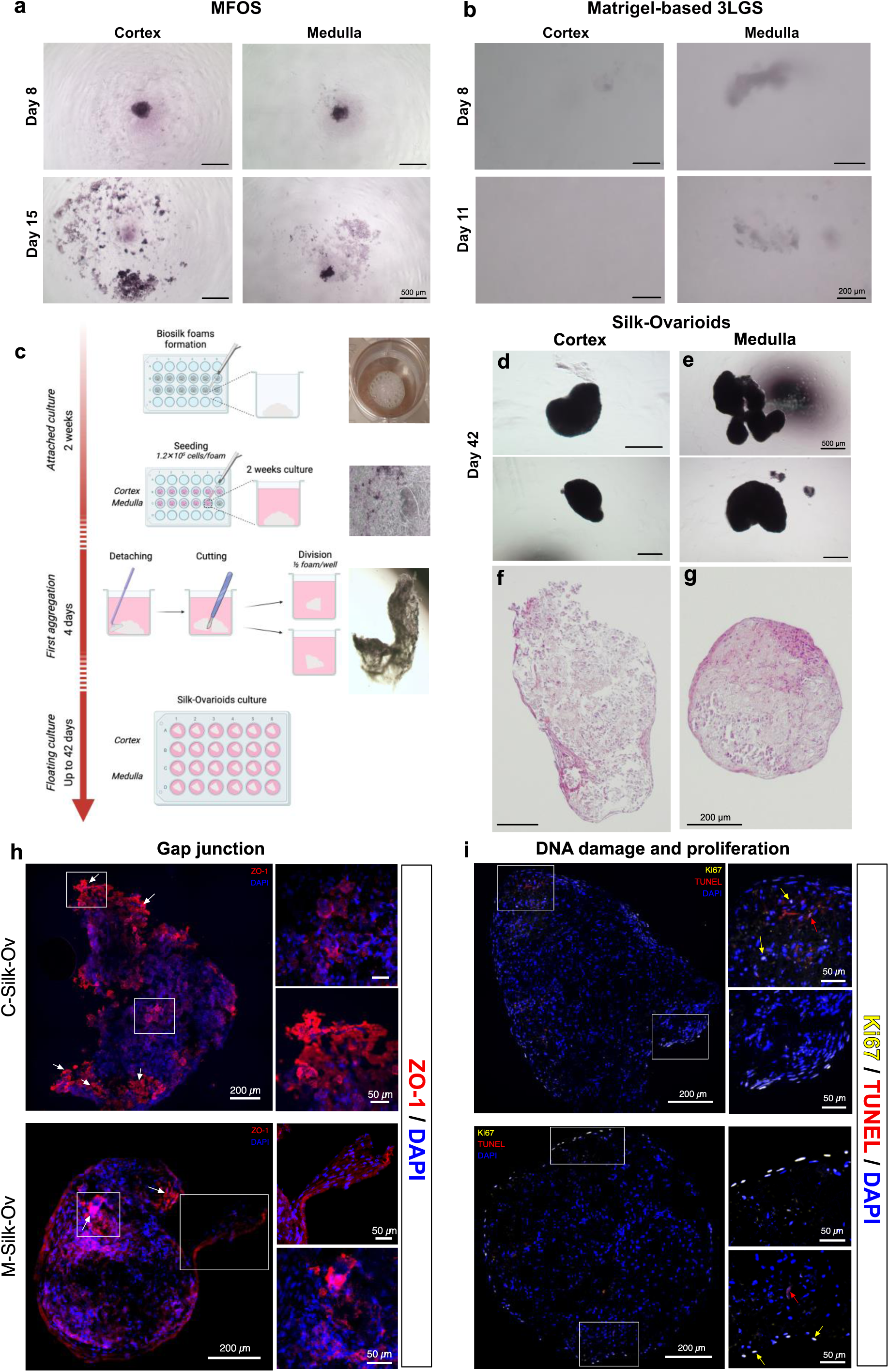
*In vitro* culture of ovarian somatic primary cells. (**a**) Representative images of MFOS of cortex and medulla in culture. Scale bar represents 500 µm. (**b**) Representative images of Matrigel-based 3LGS in culture. Scale bar represents 200 µm. (**c**) Schematic method description of Silk-Ovarioid formation and culture with the corresponding macroscopic images and timeline (Created with BioRender.com). Macroscopic morphology of (**d**) C-Silk-Ov and (**e**) M-Silk-Ov. Scale bar represents 500 µm. H&E-stained section of (**f**) C-Silk-Ov and (**g**) M-Silk-Ov. Scale bar indicates 200 µm. Representative images of immunofluorescent staining of gap junction marker ZO-1 (**h**), proliferation marker Ki67 (yellow) and DNA damage (TUNEL staining, red) (**i**) in C-Silk-Ov and M-Silk-Ov samples. Arrows of different colors indicate Biosilk background (white), Ki67 (yellow) and the TUNEL (red) fluorescence. Scale bar in the large image represents 200 µm while scale bar in inserts indicates 50 µm. All images were brightness- and contrast-adjusted to better visualize the signals. 3LGS, 3-layer gradient system; C-Silk-Ov, Cortex-derived Silk-Ovarioids; MFOS, matrix-free ovarian spheroids; M-Silk-Ov, Medulla-derived Silk-Ovarioids; TUNEL, terminal deoxynucleotidyl transferase dUTP nick end labeling; ZO-1, zona occludens-1.

Lastly, ovarian primary cells were seeded on Biosilk scaffolds (Fig. 2c). The seeded scaffolds were left untouched for two weeks, still attached to the bottom of the well, and were then detached and lifted to promote floating cell-driven aggregation. After four days of floating culture, the seeded silk foams started to compact, and they could be kept in culture for up to 42 days when we harvested them for further analyses (Fig. 2d,e). This process was performed for each patient-specific Silk-Ovarioid culture, considered a separate individual batch (n = 24 Silk-Ovarioids *per* batch) (Suppl. Table S1). Well-defined structures (400-1000 µm) could be observed under an inverted phase contrast microscope (Fig. 2d,e) and by hematoxylin and eosin (H&E) staining, which confirmed the presence of intact cells throughout the whole structure (Fig. 2f,g). Notably, a negative control, *i.e.*, cell-free Silk aggregation, was included in the first experiment and remained as a filamentous scaffold even after two weeks of floating culture. A total of five batches of culture using five patients were carried out with nearly 100% success (Suppl. Table S1).

To ascertain the quality of the Silk-Ovarioids, we detected *de novo* formation of gap junctions (zona occludens-1, ZO-1; Fig. 2h) and DNA fragmentation/proliferation (terminal deoxynucleotidyl transferase dUTP nick end labeling, TUNEL, and Ki67; Fig. 2i). Overall, the Silk-Ovarioids showed the presence of gap junctions, a low percentage of apoptotic cells – as also confirmed by the absence of the DNA damage marker, γ-H2A.X, and the cleaved form of caspase 3 (Suppl. Fig. S1) – and proliferative cells mainly present on the external part of the structure (Fig. 2h, i).

In summary, the MFOS and Matrigel-based 3LGS culture systems could not support human ovarian somatic cell growth in vitro. In contrast, the cells readily grew in Silk-Ovarioids, forming large aggregates that could be maintained for over a month in culture (Suppl. Table S1).

### Silk-Ovarioids harbor all main ovarian cell populations

To characterize the identity of the cells constituting the Silk-Ovarioids, we analyzed the presence of cell-specific markers at both RNA and protein levels. Based on previous single-cell RNA-seq (scRNA-seq) data of ovarian tissue (Wagner *et al*, 2020), we selected six markers that are highly specific to different cell types: *PDGFRA* (stromal cells), *AMHR2* (granulosa cells), *GPIHBP1*, and *CLDN5* (endothelial cells), and *MCAM* and *GJA4* (perivascular cells) (Fig. 3a, Suppl. Fig. S2). The expression of these markers was analyzed in cortex-(n=5) and medulla-derived (n=5) Silk-Ovarioids through RNA-fluorescence *in situ* hybridization (RNA-FISH) and/or immunofluorescence staining (Fig. 3b,c).

**Figure 3.**
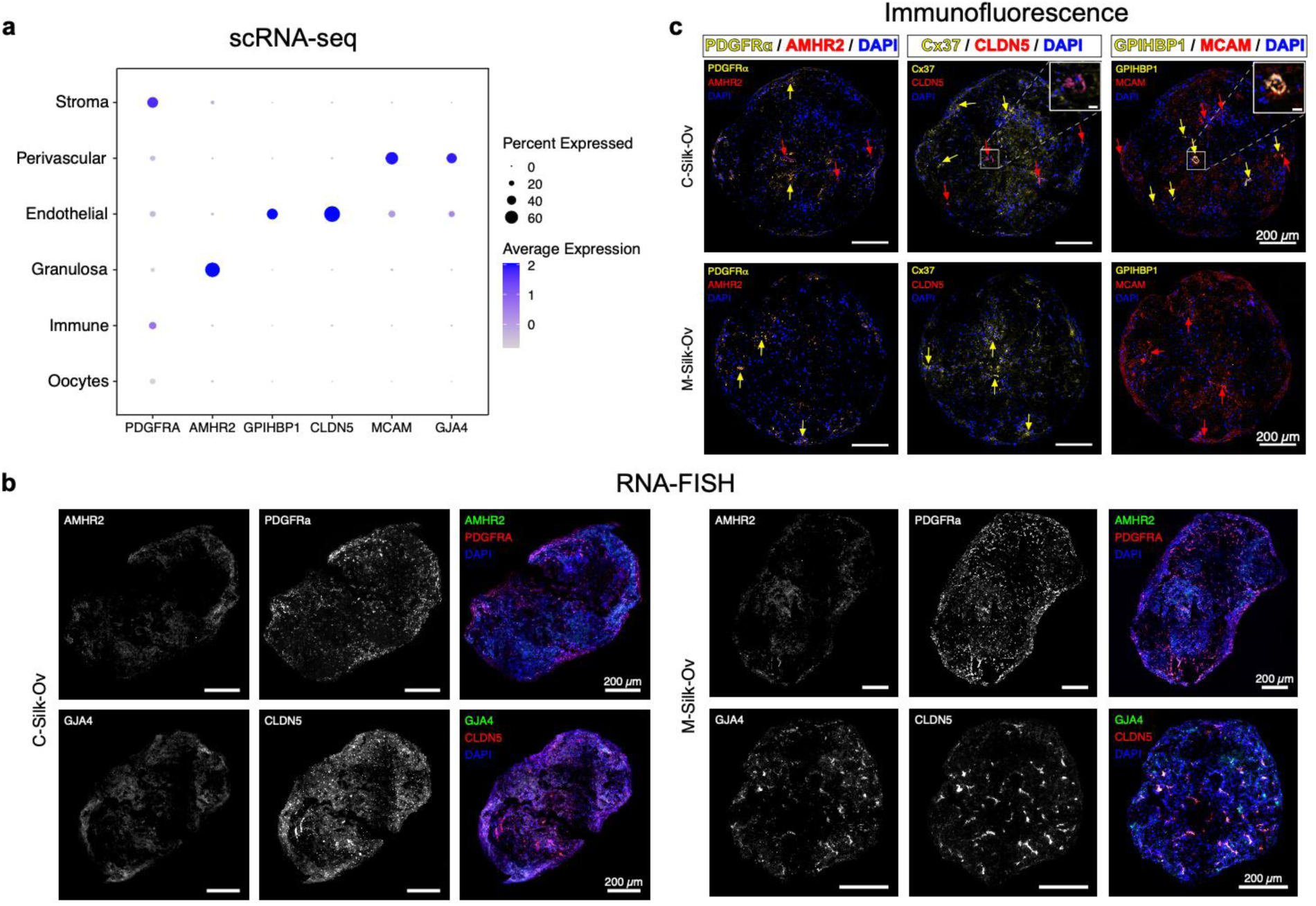
RNA and protein expression of cell type markers in Silk-Ovarioids. (**a**) Expression of selected markers (*PDGFRA, AMHR2, GPIHBP1, CLDN5, MCAM,* and *GJA4*) for stromal, perivascular, endothelial, and granulosa cells in a previously published ovarian scRNA-seq data (Wagner *et al*, 2020). Color represents the average expression in each cell type, while the size of the dot indicates the percent of cells expressing the marker. (**b**) RNA fluorescence *in situ* hybridization of stromal (*PDGFRA*), granulosa (*AMHR2*), endothelial (*CLDN5*), and perivascular (*GJA4*) cellular markers in C-Silk-Ov and M-Silk-Ov samples (n=5 each). Scale bar represents 200 µm. (**c**) Protein expression of selected markers in C-Silk-Ov and M-Silk-Ov in consecutive sections. Additional endothelial (GPIHBP1) and perivascular (MCAM) markers were used to confirm the localization of these two cell types. Red and yellow arrows pointed at the specific marker expression in the C- and M-Silk-Ov. Scale bar in the large image represents 200 µm while scale bar in inserts indicates 20 µm. All images were brightness- and contrast-adjusted to better visualize the signals. AMHR2, anti-Mullerian hormone receptor type 2; CLDN5, claudin 5; C-Silk-Ov, Cortex-derived Silk-Ovarioids; GJA4, Gap junction α 4 (a.k.a. protein connexin 37, Cx37); GPIHBP1, glycosylphosphatidylinositol-anchored high-density lipoprotein-binding protein 1; MCAM, melanoma cell adhesion molecule; M-Silk-Ov, Medulla-derived Silk-Ovarioids; PDGFR-A or -α, platelet-derived growth factor receptor α.

As shown in Fig. 3b and c, all the markers were detected in the Silk-Ovarioid samples, both at the mRNA and protein levels. All Silk-Ovarioids showed a high mRNA signal of *PDGFRA* (Fig. 3b), while the protein product was less evident (Fig. 3c). Only a few signals for AMHR2 (on both mRNA and protein levels) were detected, indicating that granulosa cells are present in very low numbers. Both endothelial (GPIHBP1 and CLDN5) and perivascular cell (MCAM and GJA4, also known as Connexin 37, Cx37) markers were found in the inner and outer part of Silk-Ovarioids and were localized near each other (Fig. 3b,c). Finally, no oocytes were detected in the heterogeneous population of somatic cells forming the Silk-Ovarioids. This was not surprising as the cortex and medulla cell suspensions were filtered with cell strainers before the cultures were started.

Notably, after six weeks of culture, the development of tubular structures in the core of Silk-Ovarioids derived from the cortex was observed. These structures were positive for the endothelial cell markers CLDN5 and GPIHBP1, suggestive of blood vessel-like formation (Fig. 3c, upper panel inserts).

### Silk-Ovarioids culture upregulates angiogenesis-related genes

To further characterize the Silk-Ovarioid model, we performed RNA sequencing on these samples (n=5 for cortex; n=6 for medulla) and compared them to ovarian tissue (n=5) and 2D cell cultures (n=4) from the same patient and cultured for the same time period alongside Silk-Ovarioids. Transcriptomic profiling of freshly collected ovarian tissues, 2D cultured cells, and Silk-Ovarioids showed a clear separation between fresh tissues and cultured samples in both cortex and medulla, explaining 68.1% and 59.4% of the difference in PC1 in principal component analysis (PCA), respectively (Fig. 4a,b). As expected, the largest number of differentially expressed genes (DEGs) was identified when tissue was compared to cultured samples (Cortex: tissue *vs* 2D culture, 2,808 DEGs; *vs* Silk-Ovarioids, 2,533 DEGs. Medulla: tissue *vs* 2D culture, 2,494 DEGs; *vs* Silk-Ovarioids, 2,246 DEGs) using a strict cutoff of FDR < 0.05, absolute log_2_ fold change > 2, and average expression > 100 (Suppl. Fig. S3a; Suppl. Table S2). The heat maps of the top 500 most variable genes confirmed a clear separation among the three types of samples (Suppl. Fig. S3b). On the other hand, comparing Silk-Ovarioids to 2D samples revealed relatively subtle effects in both cortex and medulla, where 553 and 618 DEGs were found with the same cutoff criteria, respectively (Suppl. Fig. S3a).

**Figure 4.**
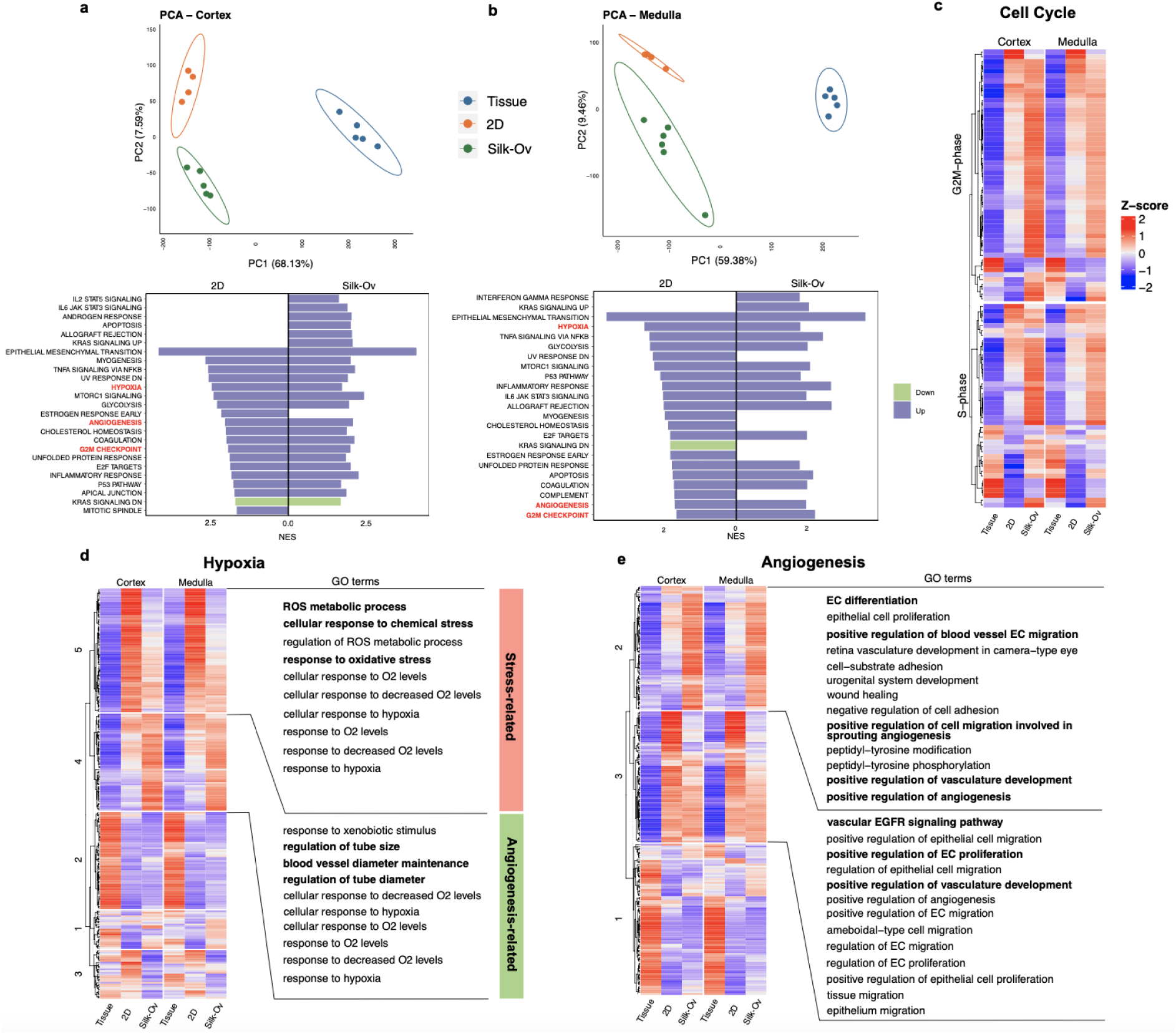
Transcriptomics profiling of tissue, 2D cell culture, and Silk-Ovarioid (Silk-Ov) samples. (**a**) PCA of tissue (n=5), 2D cell culture (n=4) and Silk-Ov (n=5 for cortex; n=6 for medulla) samples originating from ovarian cortex primary cells from 5 patients. Enriched hallmark gene sets using DEGs from 2D *vs* tissue and Silk-Ov *vs* tissue comparison. One Silk-Ov sample from cortex was removed due to low library size. (**b**) PCA of tissue (n=5), 2D (n=4) and Silk-Ov (n=6) samples originated from ovarian medulla primary cells from 5 patients. Enriched hallmark gene sets using DEGs from 2D *vs* tissue and Silk-Ov *vs* tissue comparison. Normalized enrichment score (NES) is presented in x-axis. Purple represents upregulation while light green represents downregulation. (**c**) Heatmap of average z-score of G2M and S phase marker genes in cortex and medulla samples. (**d**) Heatmap of average z-score of genes related to hypoxia in cortex and medulla samples (left). Genes are clustered using k-means clustering. Top 10 GOs enriched in cluster 5 and cluster 4 are shown (right). Stress-related cluster is shown in pink while angiogenesis-related cluster is presented in green. (**e**) Heatmap of average z-score of genes related to angiogenesis in cortex and medulla samples (left). Genes are clustered using k-means clustering. Top GOs enriched in cluster 2 and cluster 3 are shown (right). Counts are normalized using DESeq2 normalization and scaled to obtain mean equals 0 and standard deviation equals 1. The final gene expression is represented using Z-score. EC, endothelial cells; GO, gene ontology; NES, normalized enrichment score; ROS, reactive oxygen species.

To compare the impact of different culture systems on the transcriptome compared to the fresh control, we performed gene set enrichment analysis (GSEA) against hallmark gene sets using significant DEGs ranked by log_2_ fold change. GSEA results revealed a similar change induced by culture in both 2D and Silk-Ovarioid contexts, *i.e.*, the upregulation of genes involved in the cell cycle G2M checkpoint, angiogenesis, hypoxia, and TNFα signaling *via* NF-κB pathways in both cortex and medulla (Fig. 4a,b). However, we also observed Silk-Ovarioid-specific changes such as IL6-JAK-STAT3 signaling in the cortex and interferon-γ response in medulla-derived Silk-Ovarioids (Fig. 4a,b).

Focusing on the commonly affected pathways in 2D and Silk-Ovarioids samples, *i.e.,* hypoxia and angiogenesis, we plotted the average expression of associated genes collected from the Gene Ontology (GO) database. For changes related to the cell cycle, we extracted the S- and G2M-phase genes used for cell cycle scoring in the Seurat package. Silk-Ovarioids samples appeared to have a high proliferating profile compared to that of 2D and tissue, as proven by a higher expression of G2M and S phase markers (Fig. 4c). Additionally, samples cultured in 3D conditions induced a relatively lower hypoxia response compared to that of 2D as shown in the gene set enrichment score plot (Fig. 4a,b).

Using a classical machine learning method, k-means clustering, we grouped hypoxia-related genes into 5 main clusters (Fig. 4d). Two identified patterns were culture-specific, where the hypoxia-5 cluster genes were highly expressed in 2D samples, and the hypoxia-4 cluster was upregulated in Silk-Ovarioids samples (Fig. 4d). We further investigated the biological processes associated with these genes using a GO over-representation analysis. Genes in the hypoxia-5 cluster indicated significant enrichment for stress-related GOs in 2D cultures, such as reactive oxygen species metabolic process and response to oxidative stress (Fig. 4d; Suppl. Table S3). On the other hand, genes in the hypoxia-4 cluster suggested enrichment of angiogenesis-related GOs in the Silk-Ovarioids, *i.e.*, regulation of endothelial tube size, diameter, and blood vessel diameter maintenance (Fig. 4d; Suppl. Table S3).

We next characterized changes related to angiogenesis signaling. Genes that were highly expressed in Silk-Ovarioids samples were associated with endothelial cell differentiation, regulation of blood vessel endothelial cell migration, and sprouting angiogenesis (Fig. 4e; Suppl. Table S4). In the 2D-specific cluster, genes were related to vascular EGFR signaling and regulation of endothelial cells (Fig. 4e; Suppl. Table S4). These observations were further confirmed by GO enrichment analysis on significant DEGs, ranked by log_2_ fold change, identified in the Silk-Ovarioid and 2D comparisons. In line with these results, GOs related to angiogenesis (*i.e.*, vasculature and blood vessel development, sprouting angiogenesis) were significantly induced in cortex-derived Silk-Ovarioids samples. Similarly, endothelium development and sprouting angiogenesis were significantly upregulated in Silk-Ovarioid culture compared to 2D samples derived from the medulla (Suppl. Table. S5).

Collectively, these findings suggest that both cortex and medulla-derived Silk-Ovarioids have a higher cellular proliferation potential at the transcript level compared to the respective 2D system. Moreover, culture-induced hypoxia markers in both 2D and 3D models, with the main difference being the stress-related hypoxia markers upregulated in the 2D model, while in Silk-Ovarioids, pro-angiogenic mechanisms dominated.

### Silk-Ovarioids culture enables de novo-angiogenesis through formation of hypoxic environment

Due to the consistent representative signatures found among biological replicates during transcriptomics analysis, we decided to further profile single Silk-Ovarioids derived from both the cortex and medulla. Transcriptomic changes were confirmed at the protein level. In total, 3 Silk-Ovarioids and the corresponding freshly fixed tissues from both the cortex and medulla from 3 different patients were subjected to individual proteomic analysis using Liquid Chromatography-Tandem Mass Spectrometry (LC-MS/MS). In both tissue and Silk-Ovarioids, we detected peptides involved in biological processes mainly related to transport and signal transduction, RNA and protein metabolism, cell-cell communication, and cell organization, adhesion, and proliferation (proteomics data repository PXD048710).

Due to the relatively low number of peptides detected *per* Silk-Ovarioid, we analyzed the normalized protein contribution as their relative expression to the total amount of peptides detected for each sample. The heatmap of proteins involved in hypoxia and angiogenesis signaling identified two distinct clusters (Fig. 5a). Most of the proteins that had a high contribution in Silk-Ovarioid samples were previously identified in the transcriptomics analysis as angiogenesis-related genes (Fig. 5a). Next, we compared the protein and RNA levels of the hypoxia- and angiogenesis-related genes through Pearson correlations. We recorded moderate to strong correlations between the normalized protein contribution and normalized RNA counts both in the cortex and medulla (Fig. 5b).

**Figure 5.**
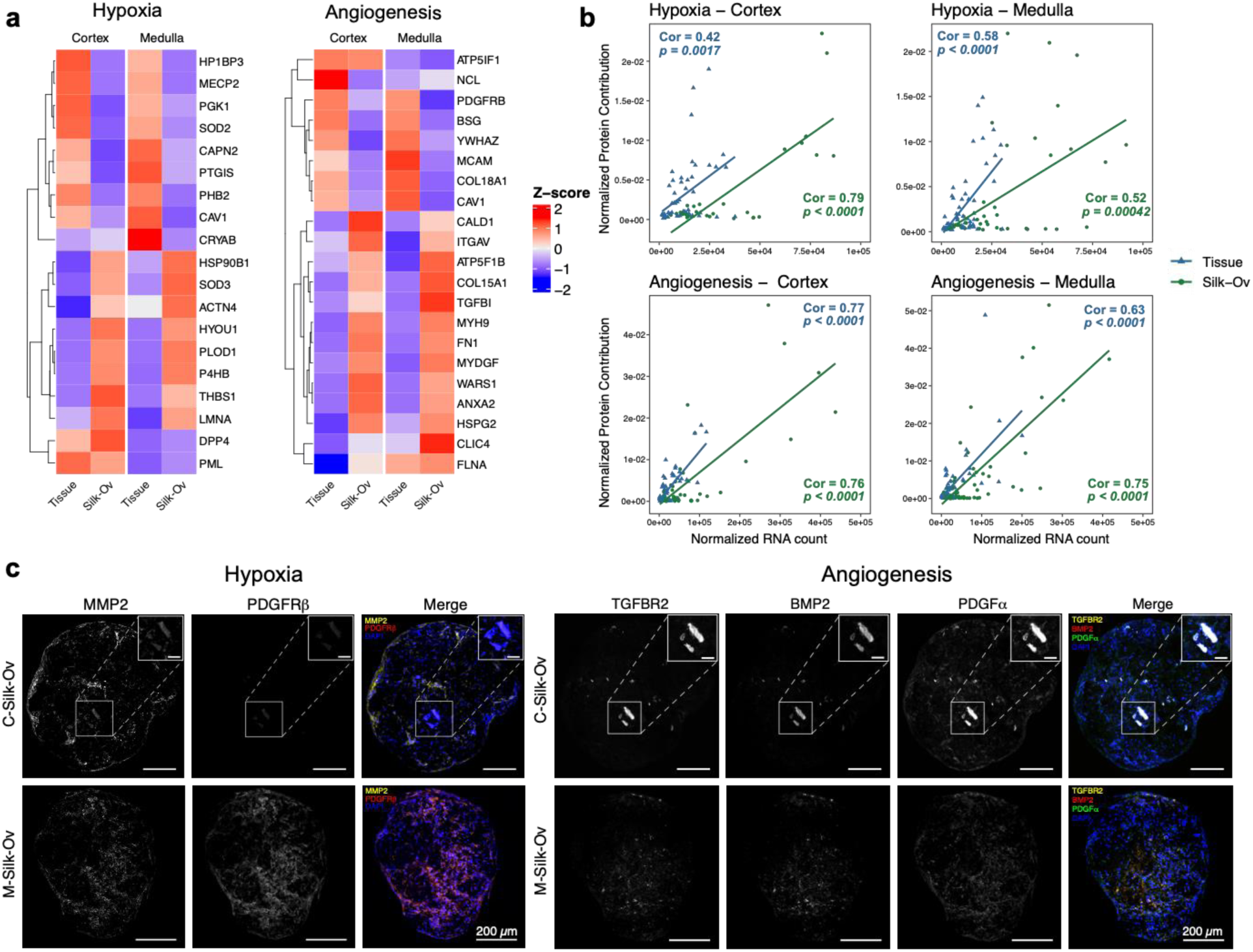
Proteomics and immunostaining of hypoxia- and angiogenesis-related markers in Silk-Ovarioids. (**a**) Average normalized protein contribution of hypoxia- and angiogenesis-related markers in tissue and C- and M-Silk-Ov samples (n=2 for cortex and n=3 for medulla). <u>One Silk-Ov sample from cortex was removed due to low concentrations of</u> <u>peptides detected.</u> Expression was normalized to obtain mean equals to 0 and standard deviation equals to 1. (**b**) Pearson correlation between detected normalized hypoxia and angiogenesis proteins and normalized RNA counts in RNA-seq data (19 markers). RNA counts were normalized using DESeq2 normalization. One Silk-Ov sample from cortex was removed due to low library size. Normalized protein contribution indicates the relative expression of a protein to the total amount of peptides detected. One Silk-Ov sample from cortex was removed due to low concentrations of peptides detected. Tissue and C- and M-Silk-Ov samples were analyzed separately. The regression lines are plotted using linear model fitting. The correlation coefficient value is shown as Cor and p value is given in the plots. Data from tissue are shown as blue triangles whilst data from Silk-Ov are presented as green round dots. (**c**) Representative images of hypoxia-(MMP2, PDGFRβ) and angiogenesis-related (TGFBR2, BMP2, PDGFα) markers immunofluorescence stainings in sequential sections of C- and M-Silk-Ov (n=5 each). Inserts show higher magnification of the vessel-like structure in the core of Silk-Ov. Scale bar in the large image represents 200 µm while scale bar in inserts indicates 50 µm. BMP2, Bone morphogenetic protein 2; C-Silk-Ov, Cortex-derived Silk-Ovarioids; MMP2, Matrix metallopeptidase 2; M-Silk-Ov, Medulla-derived Silk-Ovarioids; PDGFα, Platelet-derived growth factor α; PDGFRβ, Platelet-derived growth factor receptor β; TGFBR2, Transforming growth factor-β receptor type 2.

To further confirm the protein-RNA correlation, we selected highly expressed markers of hypoxia (*i.e.*, MMP2 and PDGFRβ) and angiogenesis (*i.e.*, TGFBR2, BMP2, and PDGFα) from the RNA-seq analysis and investigated their protein localization *via* immunofluorescent staining. The results validated the presence of hypoxic and angiogenic proteins, and their localization highlighted a functional asset of the expressed markers. In fact, the pro-angiogenic environment was surrounded and initiated by hypoxic pockets that disappeared when endothelial cells were reorganized into vessel-like structures (Fig. 5c). For instance, in Fig. 5, representative images of a cortex-derived Silk-Ovarioid already containing a vessel (Fig. 5c, upper panel), and a medulla-derived Silk-Ovarioid in which the sprouting of new vessels was still not evident (Fig. 5c, lower panel) are shown. As hypothesized, when the formation of the vessel commences, the hypoxia-related markers are downregulated in the inner part of the cortex-derived Silk-Ovarioid, while the newly created vessel showed the colocalization of all three angiogenesis-related markers (Fig. 5c, upper panel). Mirroring this pattern, in the medulla-derived Silk-Ovarioids, where the formation of the vessels was still in progress, the hypoxic markers were mainly identified in the center of the 3D model (Fig. 5c, lower panel). Concomitantly, the angiogenic markers are faintly present and localized in the core of the Silk-Ovarioid, surrounded by the pro-angiogenic hypoxic environment (Fig. 5c, lower panel).

Collectively, these observations suggest that some of the main markers driving hypoxia and angiogenesis could be detected in the proteomics analysis, positively correlating with the transcriptomic profile of both tissue and Silk-Ovarioids. Moreover, *de novo* angiogenesis could be observed in the center of the Silk-Ovarioid samples, initiated by the formation of a hypoxic environment.

### Silk-Ovarioid enables ECM formation and remodeling during culture

When comparing the transcriptomic profile of tissue and Silk-Ovarioids, the differential expression analysis and GO enrichment analysis identified a set of highly modulated targets related to extracellular matrix organization (Suppl. Table S5). Among them, we selected the most highly expressed genes, *COL1A1* and *LAMA1,* for further validation. Interestingly, we recorded a significantly higher expression of these two markers in RNA-seq data in both cortex- and medulla-derived Silk-Ovarioids compared to tissue, while at the proteomics level, only the collagen type 1 α1 (Col1α1) expression was captured and showed a similar trend to the transcriptomics data (Fig. 6a, b). The presence of *de novo* secretion of these ECM-related proteins was further confirmed by immunofluorescence staining (n=5 for both cortex- and medulla-derived Silk-Ovarioids from 5 patients), where the colocalization of both Col1α1 and laminin subunit α 1 (Lamα1) was detected in the core and outer part of the Silk-Ovarioid samples from both cortex and medulla (Fig. 6d).

**Figure 6.**
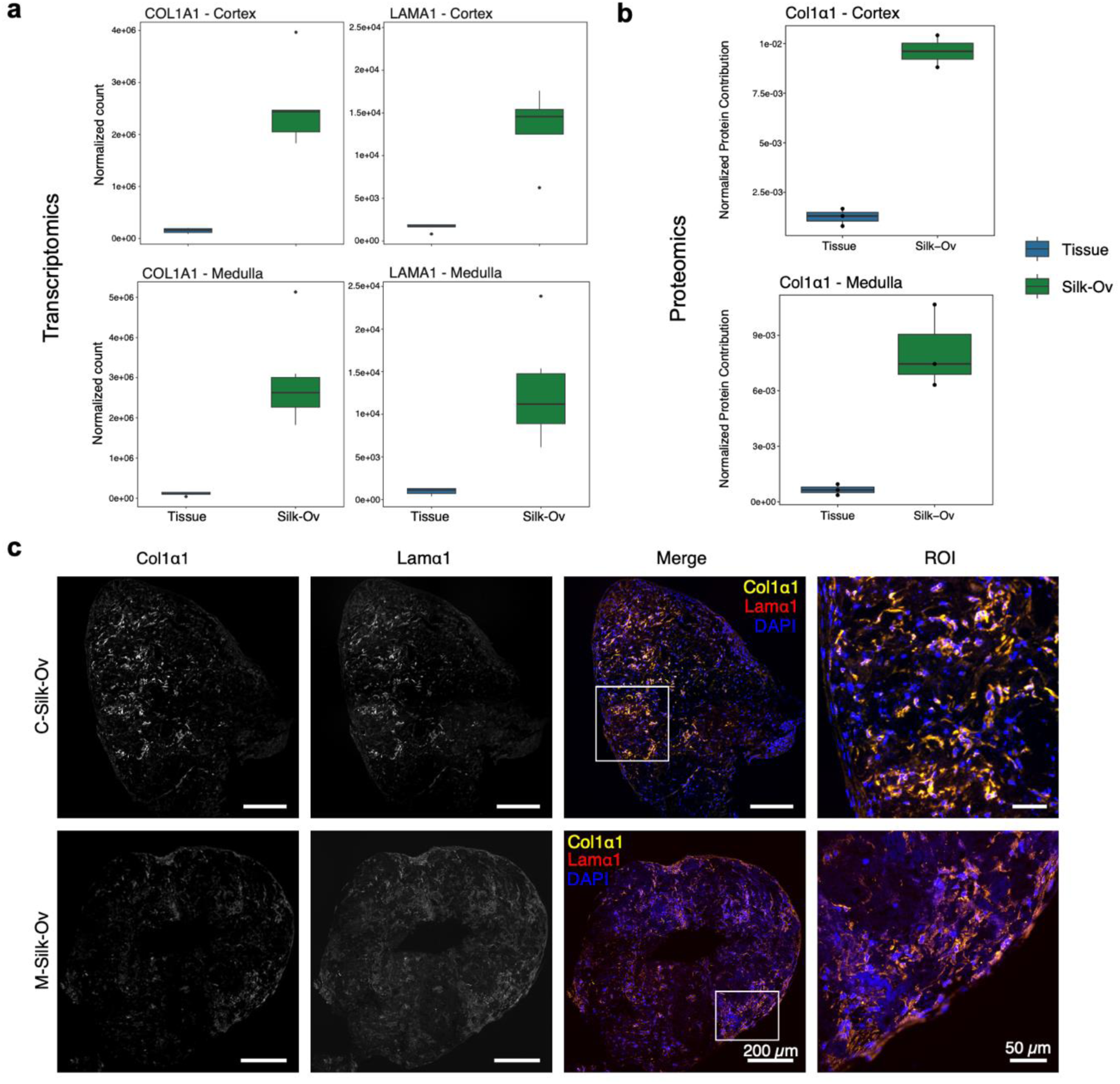
Expression of ECM markers in tissue and Silk-Ov samples in cortex and medulla. (**a**) Normalized expression of top expressed ECM related markers *COL1A1* and *LAMA1* in tissue (n=5) and Silk-Ov (n=5 for cortex; n=6 for medulla) samples in cortex and medulla in RNA-seq data. One Silk-Ov sample from cortex was removed due to low library size. RNA counts were normalized using DESeq2 normalization. (**b**) Normalized protein contribution of Col1α1 in proteomics data in tissue and Silk-Ov samples (n=3 each). Normalized protein contribution indicates the relative expression of a protein to the total amount of peptides detected. One cortex Silk-Ov sample was removed due to the low concentrations of peptides detected. Dots represent biological replicates in each group. (**c**) Representative images of Col1α1 and Lamα1 immunofluorescence staining in C-Silk-Ov and M-Silk-Ov (n=5 each). The ROIs highlight the presence of ECM markers after cell secretion. Scale bar for large images represents 200 µm while in inserts scale bar indicates 50 µm. Col1α1, Collagen type 1 α1; C-Silk-Ov, Cortex-derived Silk-Ovarioid; ECM, extracellular matrix; Lamα1, Laminin subunit α1; M-Silk-Ov, Medulla-derived Silk-Ovarioid; ROI, region of interest.

### Silk-Ovarioids secret pro-angiogenic cytokines and steroids

We conducted functional studies by measuring cytokines and steroids secreted during culture by the Silk-Ovarioids. To detect a wide range of cytokines, we used a multiplex panel covering 34 cytokines. Similar cytokines were detected in the collected medium after 42 days of culture in both cortex- and medulla-derived Silk-Ovarioids (n=4), with the most secreted being IL-6, IL-8, MCP-1, CXCL1, and SDF-1 alpha (Fig. 7a). As the cytokines were measured in culture medium, we could not compare the profiles between tissue and Silk-Ovarioids. Therefore, we examined the differences between tissue and Silk-Ovarioids samples using the RNA-seq data. Consistent with the secreted detection, the selected cytokine-encoding genes were highly expressed in Silk-Ovarioid samples compared to the cortex and medulla tissue (Fig. 7b). Interestingly, all the detected cytokines in the medium, particularly the most expressed ones (*e.g.*, IL-6, IL-8, and GM-CSF), are pro-angiogenic, stimulating endothelial cellular proliferation and migration, thus promoting the formation of new vessels under physiological conditions (Kushner *et al*, 2010).

**Figure 7.**
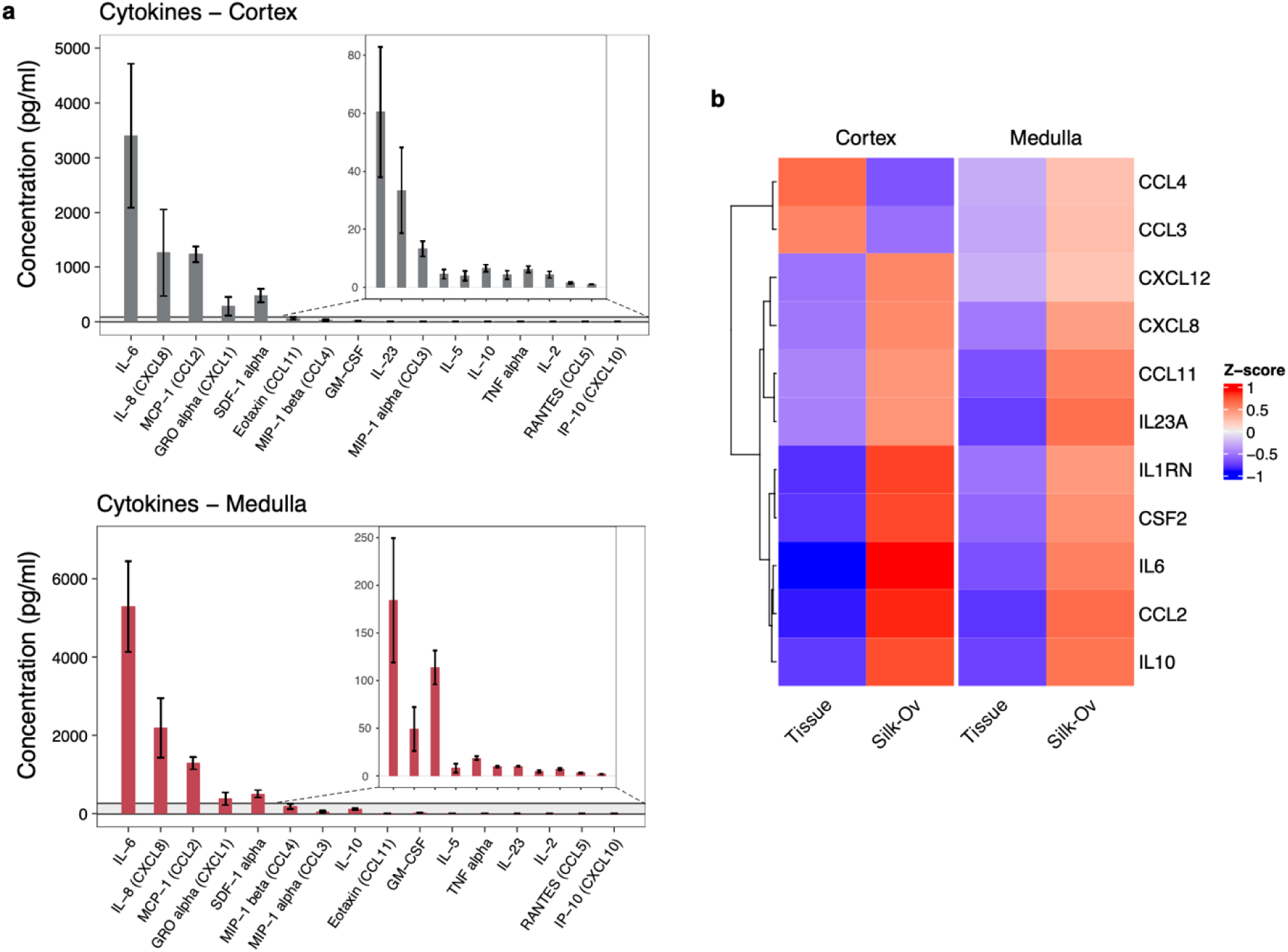
Expression and secretion of cytokines by Silk-Ovarioids. (**a**) Concentration of secreted cytokines and chemokines in the media collected from cortex and medulla Silk-Ovarioids (n=4 biological replicates and 3 technical replicates each) at day 42 of culture. Results are presented as bar plot with the mean ± SEM. (**b**) Heatmap of average z-score of genes encoding the culture medium-detected cytokines in tissue (n=5) and Silk-Ov (n=5 for cortex; n=6 for medulla) samples in cortex and medulla in RNA-seq data. One Silk-Ov sample from cortex was removed due to low library size. Counts were normalized using DESeq2 normalization and scaled to obtain mean equals 0 and standard deviation equals 1. SEM, Standard error of the mean; Silk-Ov, Silk-Ovarioid.

We identified four main steroids in the culture medium of cortex- and medulla-derived Silk-Ovarioids (n=3) after 42 days of culture. Generally, steroids were present at very low levels, often below the limit of quantification (LOQ). Pregnenolone and epitestosterone were most often above the LOQ, while the detection rate of estrogens (estrone and β-estradiol) was much lower (Suppl. Table S6). Additionally, as a general overview of the steroidogenesis-related gene expression, we compared the transcript profiles between Silk-Ovarioids and tissue. The heatmap of well-known steroidogenic enzymes and related regulators showed a downregulation of the typical granulosa (*i.e.*, *FOXL2*, *KIT*, *AMH*, *AR*) and oocyte-specific markers (*i.e.*, *GDF9*) (Suppl. Fig. S4). However, we observed an increased expression of *CYP19A1* in the Silk-Ovarioids samples, along with members of the nuclear receptor family (*e.g.*, *ESRRA*, *ESRRB*, *NR4A3*), and steroidogenesis-related growth factors and receptor genes (*e.g.*, *GNDF*, *FGF2*, *IGF2R*) (Suppl. Fig. S4). Although downregulated in Silk-Ovarioids compared to tissue samples, the presence of *CYP11A1* justifies the low but detectable secretion of pregnenolone in the culture media. Additionally, the upregulation of E1- and E2-synthesizing enzymes coding genes, *AKR1C3* and *CYP19A1,* respectively, is evidenced in Silk-Ovarioids’ transcriptomic profile compared to the tissue counterpart.

### Silk-Ovarioids as a potential new scaffold for ovary models

To better visualize the interplay between hypoxia and angiogenesis in our samples, we used the gene-concept network. Specifically, we visualized the connections of DEGs identified in the comparison of Silk-Ovarioids *vs* tissue involved in hypoxia and angiogenesis in both cortex and medulla (Fig. 8a,b). Intriguingly, crosstalk between hypoxia and angiogenesis signaling was discovered in both cortex- and medulla-derived Silk-Ovarioids, mediated by two genes, *STC1* and *VEGFA* (Fig. 8a,b). These factors, namely stanniocalcin-1 and vascular endothelial growth factor A, are not only well-known regulators of angiogenesis but also involved in follicular activation and growth (Guzmán *et al*, 2023; Bishop *et al*, 2021).

**Figure 8.**
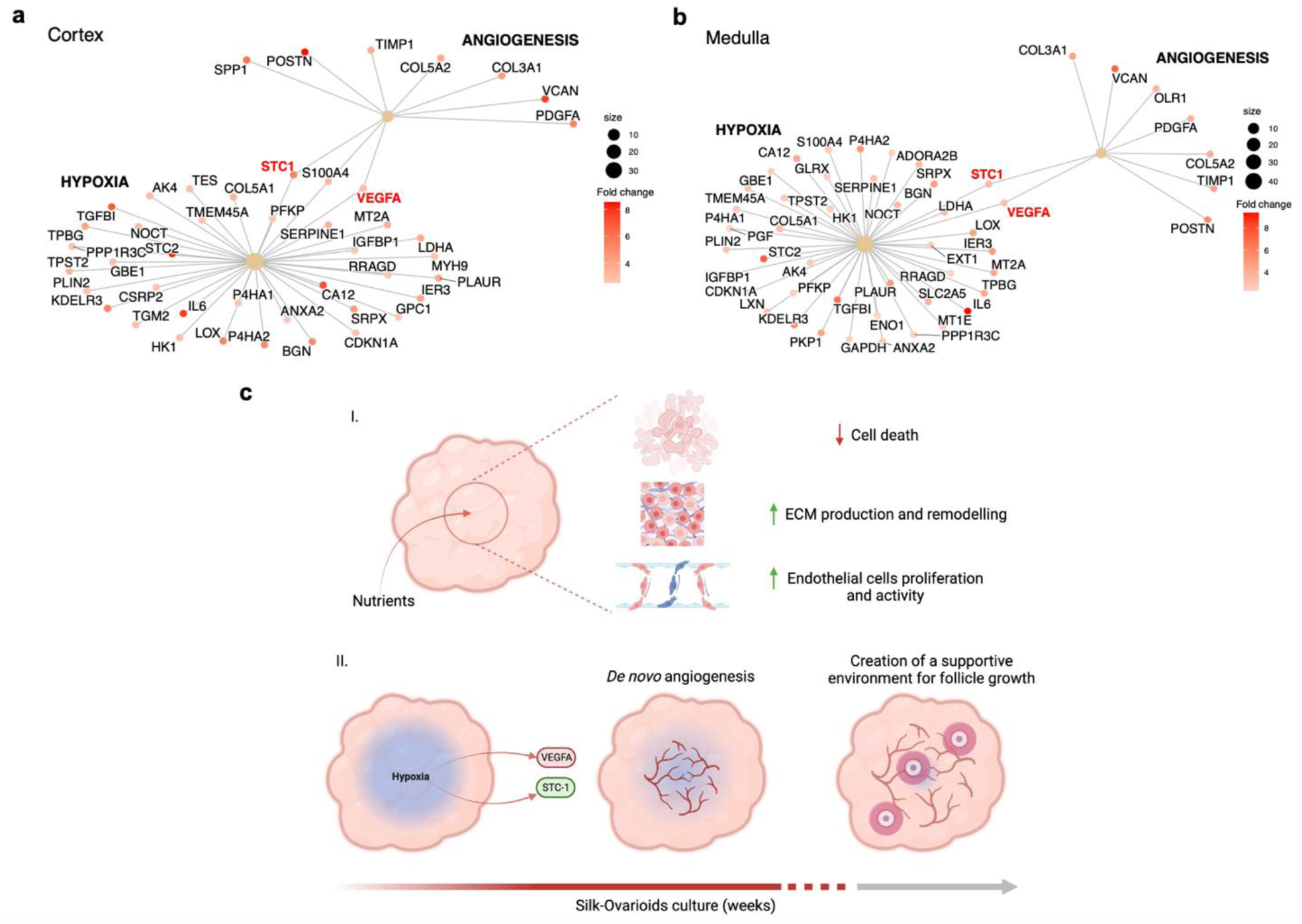
Gene-concept network and schematic overview of Silk-Ovarioids. Gene-concept network of hypoxia and angiogenesis signaling using significant DEGs identified in Silk-Ov and tissue comparison in cortex (**a**) and medulla (**b**). The color scale of the gene dots indicates fold change. The size of the central hubs’ dots (hypoxia and angiogenesis) represents the number of DEGs enriched in each gene set. (**c**) Schematic overview of (**I**) main ongoing mechanisms and (**II**) hypoxia-triggered angiogenesis process in Silk-Ov culture (Created with BioRender.com). ECM, extracellular matrix; STC-1, stanniocalcin 1; VEGFA, vascular endothelial growth factor A.

Altogether, the deep molecular characterization of the Silk-Ovarioid model showcases a promising new *in vitro* system that offers several advantages over other comparable models (Fig. 8c): the silk scaffold allows for the attachment and long-term stable growth of human adult ovarian somatic cells preserving cellular diversity; it triggers the cells to generate their own natural ECM; it promotes the passage of nutrients thus avoiding the formation of a necrotic core; and it enables the establishment of a pro-angiogenic hypoxic environment promoting proliferation and migration of endothelial cells as well as the formation of vessel-like structures (Fig. 8c-I, II). We hypothesize that the sprouting of these small channels may improve the oxygenation in the core of Silk-Ovarioids, acting as negative feedback on the hypoxic environment (Fig. 8c-ii). Furthermore, the model showed little variability between samples and patients, which will be important for possible future translation to patient-specific applications. In summary, Silk-Ovarioids can be considered as autonomous structures that, guided by the appropriate growth factors and media composition, can likely create a supportive auto-sufficient environment for *in vitro* follicular growth (Fig. 8c-ii).

## Discussion

This study introduces a novel, reproducible, and robust silk-based 3D model of primary human somatic cells derived from healthy ovaries. In contrast to previous Matrigel-based approaches for culturing patient-derived ovarian cancer cells (Nanki *et al*, 2020; Maenhoudt & Vankelecom, 2021; Yang *et al*, 2021), or testicular and ovarian organoids (Alves-Lopes *et al*, 2017; Oliver *et al*, 2021), our Silk-Ovarioids model facilitates the survival and proliferation of the main somatic cell populations of the ovary, along with the autonomous establishment of a hypoxic environment crucial for subsequent neo-angiogenesis.

The Matrigel-based 3LGS method efficiently establishes fetal gonadal organoids, supporting the maintenance of ovarian germ cells isolated from first-trimester gonads (Oliver *et al*, 2021). However, when applying the matrix-free and 3LGS organoid protocols to adult ovarian samples, the cells appeared incapable of either aggregating or surviving in culture for extended periods. A potential explanation for this protocol discrepancy is the stemness level of the adult ovary samples. Adult cells can be classified as multipotent or completely differentiated cells. In both scenarios, adult cells exhibit very low levels of stemness markers (Suppl. Table S2), or the capacity for multiple cellular divisions (Łos *et al*, 2019). This characteristic, distinct from ovarian cancers and fetal tissues, led us to employ the silk-based system, which is xeno-free and potentially compatible with future clinical applications, such as *in vitro* folliculogenesis and auto-transplantation in humans. Although the system has been applied in 3D cultures of human pluripotent stem cells (Fiorenzano *et al*, 2021; Sozzi *et al*, 2022) and several primary cell types (Johansson *et al*, 2019), it has not yet been utilized for ovarian cells. In this study, seeding ovarian primary cells on silk foams enabled their attachment and growth while still attached to the plate. Subsequently, following detachment and division of the seeded foams, all Silk-Ovarioids began to condense and aggregate after 4 days and were successfully maintained in suspension culture for up to 42 days. The condensation of the foams to create well-defined and compacted structures was solely driven by cell-to-cell interactions, as cell-free foams did not reach a compact stage even after 2 weeks of culture in suspension. This study thus represents the first description of relatively large 3D structures (ranging from 400 to 1000 µm) derived from ovarian primary cells that can be stably cultured for extended periods.

The quality of the newly established Silk-Ovarioids model was evaluated by examining their internal structures and probing the health status of the cells in terms of DNA fragmentation and proliferation. A primary concern with 3D cultures is the development of a necrotic core due to limited nutrient and oxygen exchange (Huang *et al*, 2017). In the Silk-Ovarioids, DNA fragmentation was nearly undetectable (n=5), indicating the absence of a necrotic core typically observed in structures exceeding 400 µm in diameter (36). Additionally, proliferation, as indicated by Ki67 expression, was mainly localized to the outermost thin layer of cells surrounding both cortex- and medulla-derived Silk-Ovarioids.

This 3D model establishes a hypoxic environment in the inner core, partially resembling multicellular tumor spheroids (Giverso & Preziosi, 2019). It is well-documented that a necrotic core induces hypoxia in the inner region of multicellular tumor spheroids, inhibiting cell division and prompting the surrounding cells to enter a quiescent state (Giverso & Preziosi, 2019). Thus, the cells inside tumor spheroids remain alive but non-proliferative, while proliferation continues in the outer shell (Giverso & Preziosi, 2019). In Silk-Ovarioids, the formation of a necrotic core is prevented by the silk-based scaffold, as also observed in similar systems, such as human brain organoids (Fiorenzano *et al*, 2021). However, we report the presence of the hypoxic inner environment both at the transcriptomic and proteomic levels.

In general, the formation of new gap junctions during organoid culture, concomitant with autonomous ECM secretion, indicates cell communication and proper remodeling of the scaffold. We confirmed tight cell adhesion both in the inner and outer regions of the model across 5 batches of Silk-Ovarioids, substantiated by the presence of the zona occludens-1 (ZO-1) protein. ZO-1, a marker of gap junctions, is typically expressed in epithelial and endothelial cells (Tornavaca *et al*, 2015) (Suppl. Fig. S2). Further identification of cell types in the Silk-Ovarioid model was confirmed by RNA-FISH and immunodetection in both cortex and medulla. Also, despite the minimal contribution of granulosa cells to the total cellular composition of the Silk-Ovarioids, both the presence and functionality of this pivotal cell type were confirmed with the detection of estrone and 17β-estradiol in the medium after long-term culture. The established functionality of cells was further corroborated by the production of ECM. Based on our RNA-seq data, we focused on the highly expressed proteins – namely collagen type I chain α1 and laminin subunit α1 – that are generally involved in the somatic support of follicles (Fan *et al*, 2019). Overall, the quality check of the newly established Silk-Ovarioids presents a compact 3D structure in which cells are functional and produce *de novo* ECM.

The hypoxic core of Silk-Ovarioids appears to be instrumental in establishing a pro-angiogenic environment, as supported by transcriptomic, proteomic, and validation analyses. Further support for this was found by the active secretion of pro-angiogenic cytokines into the culture media. We hypothesize that this event could mainly be initiated and perpetuated by endothelial cells in the Silk-Ovarioid model. In fact, the 3D environment allows endothelial cells to grow, exert their functions, and remodel the ECM to form vessel-like structures, as described in our study. Consistent with our results, successful co-culture of multiple cell types and the formation of micro-vessels have been shown in other silk-based models using both human and mouse-derived material (Johansson *et al*, 2019). This underscores the important role of vascularization in 3D cultures to ensure oxygen and nutrient exchange in the inner core (Loh & Choong, 2013), as well as in ovarian 3D models. In a bioprosthetic ovary created using mouse primary cells, the microporous bio-printed scaffold allowed the growth of mouse cells, the attachment of follicles, and its vascularization after grafting back to the mouse (Laronda *et al*, 2017). This process was vital not only for the survival of the grafts *per se* but also for the growth of the seeded follicles as it promoted the exchange of necessary oxygen, hormones, and nutrients for *in vivo* folliculogenesis post-transplantation (Laronda *et al*, 2017). Additionally, several studies highlighted the importance of pro-angiogenic factors (*e.g.*, VEGF, estrogen metabolites) (Henríquez *et al*, 2020; Guzmán *et al*, 2023) and cytokines (*e.g*., leukemia inhibitory factor, fibroblast growth factor, interleukins (IL) 6 and 8, and transforming growth factor β family) (Adamczak *et al*, 2021) in ovarian follicle activation, growth, and development. For instance, the downregulation of estrogen metabolites and VEGF, reported in women diagnosed with polycystic ovarian syndrome, caused the arrest of follicular growth (Henríquez *et al*, 2020). The secretion of the pro-angiogenic cytokines (*i.e.*, IL-6 and IL-8) by the Silk-Ovarioids and increased expression of *VEGFA* on a transcriptomic level are in line with the activation of angiogenic mechanisms and the presence of endothelial cells-derived tubular structures. Moreover, VEGFA, together with the hypoxia-induced STC-1, a factor involved in follicular angiogenesis as outlined in pigs (Basini *et al*, 2009) and by our gene-concept hubs, will likely establish a favorable environment for follicle activation and growth. Indeed, a recent review outlined the putative role of STC-1 in the female reproductive system of mammals, including the ovarian functions connected to insulin-like growth factor activity, steroidogenesis, and ovulation (Bishop *et al*, 2021).

## Concluding Remarks

This study demonstrates, for the first time, the successful formation of a human 3D ovarian model that can survive long-term culture without undergoing necrosis or apoptosis, maintaining healthy cells across the entire structure. This is a first step towards establishing a stable and functional ovary model that can be exploited for numerous applications, from pharmacology and toxicology to basic biology research and clinical medicine. Our Silk-Ovarioids could be utilized for *in vitro* folliculogenesis and also as transplantable ovarian grafts due to the low-immunogenic and xeno-free characteristics of the silk scaffold. One limitation, however, of this study could be that the ovarian biopsies are collected from patients undergoing androgen treatment, which raises the question of whether this could change the behavior of the cells in Silk-Ovarioids retrieved from androgen-unstimulated patients. Thus, further characterization and protocol optimization for the establishment of Silk-Ovarioids is needed to obtain a deeper understanding of this model and its future use in diverse patient groups. To create a xeno-free and defined environment for the growth of Silk-Ovarioids, fetal bovine serum (FBS) supplemented with the culture medium could be substituted by human serum albumin (HSA). Finally, the response of the Silk-Ovarioids to the stimulation from added hormones, such as follicle-stimulating hormone necessary for follicle growth, needs to be tested. This part will be achieved in the future continuation of this study by the addition of follicles to the forming Silk-Ovarioids to test their ability to support *in vitro* follicle growth and development. Overall, Silk-Ovarioids open new avenues for the development of human-based artificial ovaries. This will broaden the possibilities of gaining knowledge on ovarian pathophysiology mechanisms and translating the ovary models into clinical applications.

## Materials and Methods

### Ovarian tissue handling and dissociation into single cell suspension

Ovarian tissue samples were collected from 5 patients (age 26±5 years) who underwent gender-affirming surgery at Karolinska University Hospital and signed informed consent for medical records data collection. Patients’ data handling was performed in accordance with the European General Data Protection Regulation guidelines. All samples retrieved from the patients were pseudonymized using random codes. On the day of the surgery, samples were transported from the operation room to the laboratory in Dulbecco’s phosphate-buffered saline with calcium, magnesium, glucose, and pyruvate (ThermoFisher Scientific, USA). Upon sample receival, medulla was separated from cortex before tissue dissociation.

Dissociation of cortex and medulla was performed following our previously published protocol (Wagner *et al*, 2020). In summary, cortex and medulla were cut into pieces (smaller than 1 mm^3^) in digestion medium consisting of DMEM/F12 (ThermoFisher Scientific, USA), 2.5% heat-inactivated fetal bovine serum (HI-FBS, Gibco, Life Technology, USA), 1 mg/ml collagenase IA (Sigma Aldrich, USA), 50 μg/ml Liberase™ TM and 10 IU/ml DNase I (Roche Diagnostics, Germany). The trimmed pieces were then dissociated for maximum 50 min in a shaking water bath at 37 °C and the reaction was terminated with equal volume of termination medium (*i.e.*, DMEM/F12 supplemented with 10% HI-FBS). Subsequently, cell suspensions were centrifuged at 300 g for 5 min, followed by resuspension in culture medium (DMEM low glucose, 10% HI-FBS, 1% Penicillin/Streptomycin) and filtered through a 40 μm cell strainer (VWR, USA). The obtained single cell suspensions of ovarian primary cells were seeded either for monolayer 2D culture (1.5 × 10^6^ cells/well) in 6-well plates (Sarstedt, Germany) or 3D culture as outlined below. In all culture systems, half of the culture medium was refreshed every second day. Pictures of the aggregates were taken at every media change through an inverted microscope (Olympus CKX41, Carl Zeiss Meditec, Germany).

### 3D systems for primary ovarian cell culture

#### Matrix-free ovarian spheroids (MFOS)

After dissociation, ovarian primary cells were seeded at different densities (3 × 10^4^, 6 × 10^4^, 1.2 × 10^5^ cells/well) in 96-well ultra-low attachment plates (Corning, USA). Cells were left untouched for two days after seeding, to allow aggregation without disrupting the culture microenvironment. From the third day onwards, half of the culture media was changed every second day.

#### Matrigel-based Three-Layer Gradient System (3LGS)

For the Matrigel-based spheroids formation, the methodology for three-layer gradient system (3LGS) as described by Alves-Lopes et al (Alves-Lopes *et al*, 2018) was followed. Briefly, we diluted Matrigel (1:1; Corning, USA) in ice-cold culture medium to make the first 5 µl base drop onto the hanging cell insert (PIHT12R48, Millipore, Germany). After the solidification of the first drop in the incubator, the primary ovarian cell suspension was diluted in 3 µl of Matrigel-culture medium (1.2 × 10^5^ cells/drop) and carefully placed on top of the base drop. The Matrigel embedded cell suspension drop was left to solidify in the incubator for 15 minutes. Lastly, the embedding Matrigel drop of 8 µl was placed on top of the existing ones, to ensure the complete covering. After solidification of the last drop for 20 minutes in the incubator, the hanging cell insert was carefully placed in a 24-well plate (Starstedt, Germany) and submerged in 600 µl of prewarmed culture medium. Half of the culture medium was changed every second day.

#### Silk-based 3D culture: Silk-Ovarioids

Biosilk™ protein solution (#BS-0101, BioLamina, Sweden) was used to create the scaffolds for cell seeding according to the manufacturer instructions. The Biosilk™ fibers mainly consist of recombinant spindroins, a spider silk protein, further biofunctionalized with arginylglycylaspartic acid (RGD)-containing motif sequence. Briefly, a 20 µl drop of Biosilk™ was placed in the center of a 24-well plate (Starstedt, Germany), and air bubbles were introduced by pipetting up and down (20–22 times), followed by the establishment of a dense and compact foam of 1 cm diameter maximum. A total of 12 foams were generated in series. Ovarian primary cells dissociated from cortex and medulla were prepared as separate single cell suspension and seeded at the chosen density (1.2 × 10^5^ cells/foam, 6 foams using cortex and 6 foams using medulla derived ovarian primary cells). The cell suspension was added to the silk foam and mixed into a homogeneous solution by pipetting 5–6 times. The seeded cells were left to stabilize on the foam in the incubator for 20 min. After stabilization, pre-warmed culture medium was carefully added drop-by-drop to the cell-foam mixture. The seeded foams were left untouched for 3 days, and afterwards, half of the media was changed every second day. After 14 days, the seeded foams were lifted from the well using a spatula and divided into 2 halves. Each separated half-foam (from now on referred to as Silk-Ovarioids) was then transferred into a flat-bottom ultra-low attachment 24-well plate (Corning, USA) and kept in free floating culture for 30-42 days. Half of the media was changed every second day and collected every 7 days for further analysis. At the end of the culture period, Silk-Ovarioids were harvested in RNALater® (Invitrogen, USA), 4% methanol-free formaldehyde (ThermoFisher Scientific, USA) and snap frozen at –80 °C for further analysis. This process was performed for each patient-specific Silk-Ovarioids culture, considered as a separate individual batch.

### Samples RNA extraction and library preparation

RNA extraction of tissues stored in RNALater® and 2D cultured cells (day 42 of culture) lysed RLT buffer (Qiagen, Germany) was performed using RNeasy Mini Kit (Qiagen, Germany) according to manufacturer’s instructions. In brief, tissues were transferred to a gentleMACS^TM^ M tube (Miltenyi Biotec, Germany) containing 350 µl of RLT buffer. Tissues were fully homogenized using RNA_01.01 program provided in gentleMACS™ Dissociator (Miltenyi Biotec, Germany). The lysate was then incubated with proteinase K (600 mAU/ml, diluted 1:60 in RNase-free water; Qiagen, Germany) to digest protein. On the other hand, 2D cultured cells were harvested in 350 µl of RLT buffer. Thereafter, the lysates from both tissues and 2D cultured cells were cleaned up before DNase I (Qiagen, Germany) treatment for DNA removal.

Total RNA of Silk-Ovarioids samples (day 42 of culture) was extracted using RNeasy Micro Kit (Qiagen, Germany) according to the manufacturer’s instructions. In summary, harvested Silk-Ovarioids were resuspended in 75 µl of RLT buffer, lysed mechanically with an insulin syringe and processed for extraction with DNase I (Qiagen, Germany) treatment. Afterwards, the yield of RNA was measured with NanoPhotometer (IMPLEN, Nordic Biolabs), with subsequent determination of quality (*i.e.*, RNA integrity [RIN] value) and quantity using Agilent Bioanalyzer 2100 (Agilent Technologies, USA). Libraries were prepared in Bioinformatics and Expression Analysis (BEA) core facility at Karolinska Institute, Sweden, from samples with RIN value > 9 and A260/A280 > 1.8, using Illumina Stranded mRNA Prep Ligation protocol (Illumina, USA). Following the manufacturer’s protocol, we used 10 ng of RNA for library preparation. Subsequently, libraries were sequenced on NovaSeq6000 platform in the National Genomics Infrastructure (NGI) at SciLife Lab, Sweden.

### RNA sequencing and data analysis

STAR aligner (version 2.7.10b) was used to map the trimmed fastq files to human genome GRCh38. Subsequently, subread package (version 2.0.1) was used to align the bam files through *featureCount* function, and to annotate using Ensembl gtf file (Homo_sapiens.GRCh38.108.chr.gtf). The pipeline used for mapping and alignment was provided in a snakemake file. Differential expression analysis was performed in R (version 4.2.3) and Bioconductor through RStudio (R Core Team, 2022; RStudio Team, 2020) using DESeq2 package. Technical replicates of the samples were collapsed using *collapseReplicates* function. One Silk-Ovarioid sample from cortex was removed due to low library size. Comparisons of Silk-Ovarioids *vs* tissue, 2D cell culture *vs* tissue, and Silk-Ovarioids *vs* 2D cell culture were performed separately in cortex and medulla dataset. To obtain differentially expressed genes (DEGs), cutoff of false discovery rate (FDR) < 0.05, absolute log_2_ fold change (log_2_FC) > 2, and average expression (baseMean) > 100 were used. Affected biological pathways were identified using Gene set enrichment analysis (GSEA) against MSigDB (Liberzon *et al*, 2011) hallmark gene sets (http://www.gsea-msigdb.org/gsea/msigdb/collections.jsp) and Gene Ontology (GO) analysis through clusterProfiler (Wu *et al*, 2021; Yu *et al*, 2012), pathview (Luo & Brouwer, 2013), DOSE (Yu *et al*, 2015), and apeglm package (Zhu *et al*, 2019). ComplexHeatmap package (Gu *et al*, 2016; Gu, 2022) was used for heatmap plotting in RStudio.

Cell cycle-related genes were collected from the cell cycle scoring function in Seurat package (Hao *et al*, 2021). Genes involved in hypoxia and angiogenesis pathways were extracted from GO database. Genes related to steroidogenesis were identified through literature. Gene expression was presented as the average scaled levels from biological replicates within each group in heatmap. The genes in the heatmap were clustered using k-means clustering to identify expression patterns. Subsequently, genes in each cluster were enriched using GO over-representation analysis to identify affected signaling pathways. Gene concept network plots related to hypoxia and angiogenesis gene sets were plotted using the enriched significant DEGs through *cnetplot* function.

### Proteomics sample preparation

Tissue samples were supplemented with 12.5 µL of 8M urea, shaken vigorously and sonicated in water bath for 5 min before 50 µL of 0.2% ProteaseMAX (Promega, USA) in 10% acetonitrile (ACN) and 100 mM Tris-HCl, pH 8.5 and 1 µl of 100x protease inhibitor (Roche, Germany) were added and mixed. Following sonication in water bath for 5 min, 36.5 µL of 50 mM Tris-HCl was added and sonicated using VibraCell probe (Sonics & Materials, Inc., USA) for 40 s with on/off pulse with a 2 s interval at 20% amplitude. Lysates were spun down at 13,000 g at 4°C for 10 min and protein concentration was determined by BCA assay (Pierce, USA). A volume of lysate corresponding to 25 µg of protein was taken and supplemented with Tris-HCl buffer up to 100 µL. Proteins were reduced by adding 3.5 µL of 250 mM dithiothreitol (Sigma-Aldrich, USA) and incubated at 45°C for 37 min while shaking at 400 rpm on a block heater. Alkylation was performed with addition of 4 µL of 500 mM iodoacetamide (Sigma-Aldrich, USA) at room temperature (RT) for 30 min in the dark. Then 0.5 µg of sequencing grade modified trypsin (Promega, USA) was added to the samples and incubated for 16 h at 37°C. The digestion was stopped with 6 µL concentrated (cc.) formic acid (FA), incubating the solutions at RT for 5 min. The sample was cleaned on a C18 Hypersep plate with 40 µL bed volume (Thermo Fisher Scientific, USA), and dried using a vacuum concentrator (Eppendorf, Germany).

Silk-Ovarioids were thawed on ice and lysed with addition of 10 µL of 8M urea and sonicated in water bath for 5 min before 70 µL of 0.5M NaCl in 50 mM Tris-HCl, pH 8.5 and 0.8 µL 100x protease inhibitor (Roche Diagnostic, Germany) were added. Following sonication in water bath for 5 min, the lysates were spun down at 13,000 g at 4°C for 10 min and protein concentration was determined by BCA assay (Pierce, USA). A volume of lysate corresponding to 7.7 µg of protein was taken and supplemented with 1M urea and 438 mM NaCl in Tris-HCl buffer up to 75 µL. Proteins were reduced by adding 2.8 µL of 250 mM dithiothreitol (Sigma-Aldrich, USA) and incubated at 37°C for 45 min while shaking at 400 rpm on a block heater. Alkylation was performed with addition of 3.1 µL of 500 mM iodoacetamide (Sigma-Aldrich, USA) at RT for 30 min at 400 rpm in the dark. Proteolytic digestion was achieved by adding 0.4 µg sequencing grade modified trypsin (Promega, USA) to the samples and incubated for 16 h at 37°C. The digestion was stopped with 4.5 µL cc. FA, incubating the solutions at RT for 5 min. The sample was cleaned on a C18 Hypersep plate with 40 µL bed volume (Thermo Fisher Scientific, USA), and dried using a vacuum concentrator (Eppendorf, Germany).

Both tissue and Silk-Ovarioid samples were labeled with TMT-10plex (Thermo Fisher Scientific, USA) isobaric reagents in two sets. Peptides were solubilized in 70 µL of 50 mM triethylammonium bicarbonate and mixed with 100 µg TMT-10plex reagents in anhydrous ACN and incubated for 2 h at RT. The unreacted reagents were quenched with 6 µL of hydroxyamine for 15 min at RT. Biological samples were then combined, dried in vacuum and cleaned on C18 Hypersep plate.

### Liquid Chromatography-Tandem Mass Spectrometry data acquisition and analysis

TMT-10plex labeled peptide samples were reconstituted in solvent A (2% ACN, 0.1% FA in water) and approximately 2 µg samples was injected on a 50 cm long EASY-Spray C18 column (Thermo Fisher Scientific, USA) connected to an Ultimate 3000 nanoUPLC system (Thermo Fisher Scientific, USA) using a 90 min long gradient: 4-26% of solvent B (98% ACN, 0.1% FA) in 90 min, 26-95% in 5 min, and 95% of solvent B for 5 min at a flow rate of 300 nL/min. Mass spectra were acquired using Q Exactive HF hybrid quadrupole-Orbitrap mass spectrometer (Thermo Fisher Scientific, USA) in a range of *m/z* 375 to 1700 at a resolution of R=120,000 (at *m/z* 200) targeting 1x10^6^ ions for maximum injection time of 80 ms, followed by data-dependent higher-energy collisional dissociation (HCD) fragmentations of top 18 precursor ions with a charge state 2+ to 7+, and using a 45 s dynamic exclusion. The tandem mass spectra were acquired with a resolution of R=60,000, targeting 2x10^5^ ions for maximum injection time of 54 ms, setting quadrupole isolation width to 1.4 Th and normalized collision energy to 34%.

Acquired raw data files were analyzed using Proteome Discoverer v3.0 (Thermo Fisher Scientific, USA) with MS Amanda v2.0 search engine against human protein database (SwissProt, 20,022 entries downloaded on 9 February 2023). A maximum of two missed cleavage sites were allowed for full tryptic digestion, while setting the precursor and the fragment ion mass tolerance to 10 ppm and 0.02 Da, respectively. Carbamidomethylation of cysteine was specified as a fixed modification. Oxidation on methionine, deamidation of asparagine and glutamine, TMT6plex (+229.163 Da) of lysine and peptide N-termini were set as dynamic modifications. Initial search results were filtered with 5% FDR using Percolator node in Proteome Discoverer. Quantification was based on the reporter ion intensities.

Proteomics data analysis was performed in R through RStudio (RStudio Team, 2020). Proteins that were detected in both tissue and Silk-Ovarioids samples were kept for downstream analysis. Normalized protein contributions were calculated by dividing the abundance of protein detected by the sum of the protein abundances in the sample. One Silk-Ov sample from cortex was removed from the analysis due to low concentrations of peptides detected. Subsequently, we identified in the proteomics data the proteins that were presented in the transcriptomics heatmap and used the scaled protein contributions to plot heatmaps. Only samples that were used for both transcriptomics and proteomics analysis were included. Thereafter, a Pearson correlation analysis was performed on data excluding outliers between the normalized RNA counts and normalized protein contributions for tissue and Silk-Ovarioids samples respectively, using *cor.test* function in RStudio.

### RNA fluorescent in situ hybridization (RNA-FISH)

*In situ* hybridization on formaldehyde-fixed paraffin-embedded Silk-Ovarioids sections (4 μm) was performed using Multiplex Fluorescent Detection Kit v2 (Advanced Cell Diagnostics, USA) following the manufacturer’s protocol. In summary, Silk-Ovarioids sections were baked, deparaffinized, and rehydrated before antigen retrieval. Afterwards, hybridization with probes for human *AMHR2* (Cat. No. 490241, Advanced Cell Diagnostics, USA), *PDGFRA* (Cat. No. 604481-C2, Advanced Cell Diagnostics, USA), *GJA4* (Cat. No. 856221, Advanced Cell Diagnostics, USA), and *CLDN5* (Cat. No. 517141-C2, Advanced Cell Diagnostics, USA) were performed at 40 °C for 2 h. Probe against Ubiquitin C served as a positive control while probe detecting bacterial gene DabB was used as a negative control (Advanced Cell Diagnostics, USA). After channel development, sections were incubated with TSA vivid dye 570 and 650 (Tocris, UK). Thereafter, DAPI (Advanced Cell Diagnostics, USA) was used to stain nuclei before mounting the samples with prolong Gold anti-fade mounting medium (Life Technology, USA). Fluorescent-labelled sections were imaged using an inverted widefield Nikon microscope at Live Cell Imaging (LCI) core facility at Karolinska Institute, Sweden.

### Immunohistochemistry and immunofluorescence of Silk-Ovarioids and tissue sections

For histological evaluation, FFPE Silk-Ovarioids samples were sectioned at 4 µm and stained with hematoxylin and eosin. The quality of the structure was evaluated through presence and localization of nuclei in inner and outer part of the organoids. Stained sections were imaged with Olympus IX81 inverted microscope (Carl Zeiss Meditec, Germany).

Immunofluorescence staining was employed to study the localization of selected markers on FFPE sections of Silk-Ovarioids and tissues samples. Antigen retrieval was performed on deparaffinized and rehydrated 4 µm sections immersed in Tris/EDTA solution (Sigma-Aldrich, USA). Blocking solution was composed of 5% w/v bovine serum albumin (BSA, Sigma-Aldrich, USA), 20% normal donkey serum (Life Technology, USA), and Tris buffered saline solution. Sections were blocked for 1.5 h at RT to reduce unspecific binding. Subsequently, primary antibodies against ZO-1, Ki67, γ-H2A.X, cleaved caspase 3, PDGFRα, AMHR2, Connexin 37 (Cx37), CLDN5, MCAM, GPIHBP1, Collagen 1 α1 (Col1α1), Laminin α1 (Lamα1), MMP2, PDGFRβ, BMP2, TGFBR2, and PDGFα (Suppl. Table S6) were added to the sections for incubation at 4 °C overnight. Isotype controls were included as negative control (Suppl. Table S6). Afterwards, secondary antibodies (Suppl. Table S6) incubation was performed at RT for 2 h in the dark. Subsequently, sections were counterstained with DAPI (ThermoFisher Scientific, USA), mounted with prolong Gold anti-fade mounting medium (Life Technology, USA), and imaged using an inverted widefield Nikon microscope using 20X/0.75 air objective with 1.5x lens at LCI core facility. Images were assembled using OMERO.figure and adjusted for brightness and contrast to better visualize the signal. All images were presented on the same scale.

### TUNEL assay

DNA damage in Silk-Ovarioids was assessed using Terminal deoxynucleotidyl transferase-mediated dUTP nick-end labeling (TUNEL) assay (Fluorescence, 594 nm, Cell Signaling Technology, USA), according to the manufacturer’s instructions. In brief, sections were deparaffinized, rehydrated, and proceeded to antigen retrieval in citrate solution (Sigma Aldrich, USA). After the TUNEL reaction equilibration, positive control was prepared by treating the sections with DNase I (>3000 IU/ml, Roche Diagnostics, Germany) for 10 min at RT. Sections used as negative control were incubated with reaction buffer only. Subsequently, TUNEL reaction mix was added to the sections, followed by 2 h incubation (37 °C).

### Luminex

A bead-based multiplex immunoassay system (Multiplex kit Cytokine & Chemokine 34-Plex Human ProcartaPlex™ Panel 1A, ThermoFisher, USA) was applied to measure the secretion of cytokines and chemokines by the Silk-Ovarioids samples into the spent culture media after 42 days of culture. In brief, cell debris was removed by centrifuging the supernatants before the assay. The supernatants (50 μl) were then mixed with magnetic beads, as obtained by washing 200 μl of bead mix in a hand-held magnetic plate washer and incubated in dark at RT for 90 min with continuous shaking (500 rpm). After washing, the beads were incubated sequentially with the detection antibody mix and 50 μl of streptavidin-phycoerythrin (SAPE) solution in the dark at RT (30 min, 500 rpm). Thereafter, the washed beads were resuspended, incubated in reading buffer (120 μl) for 5 min with continuous shaking at 500 rpm before reading signals. Luminex FLEXMAP 3D instruments coupled with xPONENT 4.3 (Luminex Corporation, USA) were used for recording the median fluorescence intensity (MFI). We then converted the MFI and presented the results in pg/ml, which was calculated based on a logistic standard curve.

### Steroid hormones detection and measurement

Steroid hormones were measured in culture media after 42 days of culture of Silk-Ovarioids samples using high performance liquid chromatography coupled with tandem mass spectrometry (HPLC-MS/MS) through an EVOQ Elite Triple Quadrupole mass spectrometer (Bruker, Germany) and an Ultimate 3000 UPLC system with a DGP-3600RS dual-gradient pump (Draskau *et al*, 2019). The limit of quantification (LOQ) for each of the measured hormones is reported in Suppl. Table S6. The limit of detection and LOQ were estimated as the concentrations corresponding to three- and ten-times signal-to-noise, respectively. Data handling was done using the software MS Workstation v. 8.2.1.

### Ethics statement

Use of ovarian tissue in research was approved by Stockholm Region Ethical Review Board (Dnr. 2015/798–31/2 with amendments). Clinicians informed the patients about the study, and cortical tissue was biopsied from ovaries collected from patients undergoing gender-affirming surgery after written and signed informed consent was obtained.

## Acknowledgments

We express our gratitude to the clinicians and research nurses that assisted us in the recruitment of patients and in biopsy collection, and to the patients who contributed the study with their invaluable clinical samples. We acknowledge Jasmin Hassan, Eleftheria Maria Panagiotou, and Nikki Wallius Hiltunen for their technical assistance in handling ovarian tissue. This study was partially performed at the Live Cell Imaging Core facility/Nikon Center of Excellence, at Karolinska Institutet, Sweden, supported by the Karolinska Institute infrastructure council. We would like to thank BEA, the Bioinformatics and Expression Analysis core facility, and FENO which are supported by the board of research at the Karolinska Institute. The authors acknowledge support from the National Genomics Infrastructure in Stockholm funded by Science for Life Laboratory, the Knut and Alice Wallenberg Foundation and the Swedish Research Council, and SNIC/NAISS/Uppsala Multidisciplinary Center for Advanced Computational Science for assistance with massively parallel sequencing and access to the UPPMAX computational infrastructure. Protein identification and quantification were carried out by the Proteomics Biomedicum core facility, Karolinska Institutet (https://ki.se/en/research/proteomics-biomedicum-core-facility). We wish to thank the Biobank and Study Support at Karolinska University Hospital for their contribution including professional service and support. We would like to thank Editage (www.editage.com) for English language editing.

## Funding

Research grant from the Center for Innovative Medicine (CIMED) at Karolinska Insitutet

European Union’s Horizon 2020 research and innovation programme (project ERIN No. 952516)

Horizon Europe grant (NESTOR, grant no. 101120075) of the European Commission Swedish Research Council for Sustainable Development FORMAS (2018-02280, 2020-01621)

StratRegen Funding from Karolinska Institute, Swedish Research Council VR (grant 2020-02132)

Swedish Childhood Cancer Fund (Reference PR2017-0044, PR2020-0096) Estonian Research Council (grant PRG1076)

European Union’s H2020 project Sinfonia (No.857253) (INL research)

SbDToolBox, with reference NORTE-01-0145-FEDER-000047, supported by Norte Portugal Regional Operational Programme (NORTE 2020), under the PORTUGAL 2020 Partnership Agreement, through the European Regional Development Fund (INL research).

## Author contributions

Conceptualization: VDN, PD, AS

Methodology: VDN, TL, ZX, KP, AD, AV, FL, EAM, MP, TS, RZ

Investigation: VDN, TL

Visualization: VDN, TL

Supervision: GA, PD, AS

Funding acquisition: EAM, TS, GA, PD, AS

Writing—original draft: VDN, TL, PD

Writing—review & editing: All the authors.

## Data and materials availability

RNA-sequencing count matrix is deposited in Gene Expression Omnibus (GEO) with accession number GSE253571; the fastq files of the same transcriptomics data are currently under submission to the Federated EGA (FEGA) Sweden data repository. Single cell RNA-seq data has been deposited in the ArrayExpress database at EMBL-EBI with the accession codes ‘E-MTAb−8381’(Wagner *et al*, 2020). The mass spectrometry proteomics data have been deposited to the ProteomeXchange Consortium via the PRIDE (Perez-Riverol *et al*, 2022) partner repository with the dataset identifier PXD048710. The code used for the analysis can be found in https://github.com/tialiv/Silk-Ovarioid_project.

## References

Adamczak R, Ukleja-Sokołowska N, Lis K & Dubiel M (2021) Function of follicular cytokines: Roles played during maturation, development and implantation of embryo. Medicina (Lithuania) 57: 1251 doi:10.3390/medicina57111251 [PREPRINT]

Alves-Lopes JP, Söder O & Stukenborg JB (2017) Testicular organoid generation by a novel in vitro three-layer gradient system. Biomaterials 130: 76–89

Alves-Lopes JP, Söder O & Stukenborg JB (2018) Use of a three-layer gradient system of cells for rat testicular organoid generation. Nat Protoc 13: 248–259

Antonouli S, Di Nisio V, Messini C, Daponte A, Rajender S & Anifandis G (2023) A comprehensive review and update on human fertility cryopreservation methods and tools. Front Vet Sci 10: 1151254

Bai X & Wang S (2022) Signaling pathway intervention in premature ovarian failure. Front Med (Lausanne) 9: 999550

Barbato V, Genovese V, De Gregorio V, Di Nardo M, Travaglione A, De Napoli L, Fragomeni G, Zanetti EM, Adiga SK, Mondrone G, et al (2023) Dynamic in vitro culture of bovine and human ovarian tissue enhances follicle progression and health. Sci Rep 13: 11773

Basini G, Bussolati S, Santini SE & Grasselli F (2009) Stanniocalcin, a potential ovarian angiogenesis regulator, does not affect endothelial cell apoptosis. Ann N Y Acad Sci 1171: 94–99

Bishop A, Cartwright JE & Whitley GS (2021) Stanniocalcin-1 in the female reproductive system and pregnancy. Hum Reprod Update 27: 1098–1114

Desai N, Abdelhafez F, Calabro A & Falcone T (2012) Three dimensional culture of fresh and vitrified mouse pre-antral follicles in a hyaluronan-based hydrogel: a preliminary investigation of a novel biomaterial for in vitro follicle maturation. Reproductive Biology and Endocrinology 10: 29

Diáz-Garciá C, Herraiz S, Such E, Andrés MDM, Villamón E, Mayordomo-Aranda E, Cervera J V., Sanz MA & Pellicer A (2019) Dexamethasone does not prevent malignant cell reintroduction in leukemia patients undergoing ovarian transplant: Risk assessment of leukemic cell transmission by a xenograft model. Human Reproduction 34: 1485–1493

Donnez J, Martinez-Madrid B, Jadoul P, Van Langendonckt A, Demylle D & Dolmans MM (2006) Ovarian tissue cryopreservation and transplantation: A review. Hum Reprod Update 12: 519–535

Draskau MK, Boberg J, Taxvig C, Pedersen M, Frandsen HL, Christiansen S & Svingen T (2019) In vitro and in vivo endocrine disrupting effects of the azole fungicides triticonazole and flusilazole. Environmental Pollution 255: 113309

Esencan E, Beroukhim G & Seifer DB (2022) Age-related changes in Folliculogenesis and potential modifiers to improve fertility outcomes -A narrative review. Reproductive Biology and Endocrinology 20: 156

Fan X, Bialecka M, Moustakas I, Lam E, Torrens-Juaneda V, Borggreven N V., Trouw L, Louwe LA, Pilgram GSK, Mei H, et al (2019) Single-cell reconstruction of follicular remodeling in the human adult ovary. Nat Commun 10: 3164

Fiorenzano A, Sozzi E, Birtele M, Kajtez J, Giacomoni J, Nilsson F, Bruzelius A, Sharma Y, Zhang Y, Mattsson B, et al (2021) Single-cell transcriptomics captures features of human midbrain development and dopamine neuron diversity in brain organoids. Nat Commun 12: 7302

Giverso C & Preziosi L (2019) Influence of the mechanical properties of the necrotic core on the growth and remodelling of tumour spheroids. Int J Non Linear Mech 108: 20–32

Graham O, Rodriguez J, van Biljon L, Fashemi B, Graham E, Fuh K, Khabele D & Mullen M (2023) Generation and Culturing of High-Grade Serous Ovarian Cancer Patient-Derived Organoids. Journal of Visualized Experiments 2023: e64878

Grosbois J, Bailie EC, Kelsey TW, Anderson RA & Telfer EE (2023) Spatio-temporal remodelling of the composition and architecture of the human ovarian cortical extracellular matrix during in vitro culture. Human Reproduction 38: 444–458

Gu Z (2022) Complex heatmap visualization. iMeta 1: e43

Gu Z, Eils R & Schlesner M (2016) Complex heatmaps reveal patterns and correlations in multidimensional genomic data. Bioinformatics 32: 2847–2849

Guzmán A, Hernández-Coronado CG, Gutiérrez CG & Rosales-Torres AM (2023) The vascular endothelial growth factor (VEGF) system as a key regulator of ovarian follicle angiogenesis and growth. Mol Reprod Dev 90: 201–217

Hao J, Li T, Heinzelmann M, Moussaud-Lamodière E, Lebre F, Krjutškov K, Damdimopoulos A, Arnelo C, Pettersson K, Alfaro-Moreno E, et al (2024) Effects of chemical in vitro activation versus fragmentation on human ovarian tissue and follicle growth in culture. Hum Reprod Open 2024: hoae028

Hao J, Tuck AR, Prakash CR, Damdimopoulos A, Sjödin MOD, Lindberg J, Niklasson B, Pettersson K, Hovatta O & Damdimopoulou P (2020) Culture of human ovarian tissue in xeno-free conditions using laminin components of the human ovarian extracellular matrix. J Assist Reprod Genet 37: 2137–2150

Hao Y, Hao S, Andersen-Nissen E, Mauck WM, Zheng S, Butler A, Lee MJ, Wilk AJ, Darby C, Zager M, et al (2021) Integrated analysis of multimodal single-cell data. Cell 184: 3573–3587.e29

Henríquez S, Kohen P, Xu X, Villarroel C, Muñoz A, Godoy A, Strauss JF & Devoto L (2020) Significance of pro-angiogenic estrogen metabolites in normal follicular development and follicular growth arrest in polycystic ovary syndrome. Human Reproduction 35: 1655–1665

Hovatta O, Silye R, Abir R, Krausz T & Winston RML (1997) Extracellular matrix improves survival of both stored and fresh human primordial and primary ovarian follicles in long-term culture. Human Reproduction 12: 1032–1036

Hu B, Wang R, Wu D, Long R, Ruan J, Jin L, Ma D, Sun C & Liao S (2023) Prospects for fertility preservation: the ovarian organ function reconstruction techniques for oogenesis, growth and maturation in vitro. Front Physiol 14: 1177443

Huang Y, Wang S, Guo Q, Kessel S, Rubinoff I, Chan LLY, Li P, Liu Y, Qiu J & Zhou C (2017) Optical coherence tomography detects necrotic regions and volumetrically quantifies multicellular tumor spheroids. Cancer Res 77: 6011–6020

Johansson U, Widhe M, Shalaly ND, Arregui IL, Nilebäck L, Tasiopoulos CP, Åstrand C, Berggren PO, Gasser C & Hedhammar M (2019) Assembly of functionalized silk together with cells to obtain proliferative 3D cultures integrated in a network of ECM-like microfibers. Sci Rep 9: 6291

Kushner E, Van Guilder G, MacEneaney O, Greiner J, Cech J, Stauffer B & DeSouza C (2010) Ageing and endothelial progenitor cell release of proangiogenic cytokines. Age Ageing 39: 268–272

Laronda MM, Rutz AL, Xiao S, Whelan KA, Duncan FE, Roth EW, Woodruff TK & Shah RN (2017) A bioprosthetic ovary created using 3D printed microporous scaffolds restores ovarian function in sterilized mice. Nat Commun 8: 15261

Liberzon A, Subramanian A, Pinchback R, Thorvaldsdóttir H, Tamayo P & Mesirov JP (2011) Molecular signatures database (MSigDB) 3.0. Bioinformatics 27: 1739–1740

Loh QL & Choong C (2013) Three-dimensional scaffolds for tissue engineering applications: Role of porosity and pore size. Tissue Eng Part B Rev 19: 485–502

Łos MJ, Skubis A & Ghavami S (2019) Stem Cells. Stem Cells and Biomaterials for Regenerative Medicine: 5–16

Luo W & Brouwer C (2013) Pathview: An R/Bioconductor package for pathway-based data integration and visualization. Bioinformatics 29: 1830–1831

Maenhoudt N & Vankelecom H (2021) Protocol for establishing organoids from human ovarian cancer biopsies. STAR Protoc 2: 100429

La Marca A, Papaleo E, D’Ippolito G, Grisendi V, Argento C & Volpe A (2012) The ovarian follicular pool and reproductive outcome in women. Gynecological Endocrinology 28: 166–169

Mehedintu C, Frincu F, Carp-Veliscu A, Barac R, Badiu DC, Zgura A, Cirstoiu M, Bratila E & Plotogea M (2021) A warning call for fertility preservation methods for women undergoing gonadotoxic cancer treatment. Medicina (Lithuania) 57: 1340

Nanki Y, Chiyoda T, Hirasawa A, Ookubo A, Itoh M, Ueno M, Akahane T, Kameyama K, Yamagami W, Kataoka F, et al (2020) Patient-derived ovarian cancer organoids capture the genomic profiles of primary tumours applicable for drug sensitivity and resistance testing. Sci Rep 10: 12581

Oliver E, Alves-Lopes JP, Harteveld F, Mitchell RT, Åkesson E, Söder O & Stukenborg JB (2021) Self-organising human gonads generated by a Matrigel-based gradient system. BMC Biol 19: 212

Ouni E, Nedbal V, Da Pian M, Cao H, Haas KT, Peaucelle A, Van Kerk O, Herinckx G, Marbaix E, Dolmans MM, et al (2022) Proteome-wide and matrisome-specific atlas of the human ovary computes fertility biomarker candidates and open the way for precision oncofertility. Matrix Biology 109: 91–120

Ouni E, Peaucelle A, Haas KT, Van Kerk O, Dolmans MM, Tuuri T, Otala M & Amorim CA (2021) A blueprint of the topology and mechanics of the human ovary for next-generation bioengineering and diagnosis. Nat Commun 12: 5603

Perez-Riverol Y, Bai J, Bandla C, García-Seisdedos D, Hewapathirana S, Kamatchinathan S, Kundu DJ, Prakash A, Frericks-Zipper A, Eisenacher M, et al (2022) The PRIDE database resources in 2022: A hub for mass spectrometry-based proteomics evidences. Nucleic Acids Res 50: D543–D552

R Core Team (2022) R: A language and environment for statistical computing. R Foundation for Statistical Computing, Vienna, Austria. https://wwwR-project.org/ 4

RStudio Team (2020) RStudio: Integrated Development for R. RStudio, PBC, Boston, MA. http://www.rstudio.com/

Shikanov A, Smith RM, Xu M, Woodruff TK & Shea LD (2011) Hydrogel network design using multifunctional macromers to coordinate tissue maturation in ovarian follicle culture. Biomaterials 32: 2524–2531

Simon LE, Rajendra Kumar T & Duncan FE (2020) In vitro ovarian follicle growth: A comprehensive analysis of key protocol variables. Biol Reprod 103: 455–470

Sozzi E, Kajtez J, Bruzelius A, Wesseler MF, Nilsson F, Birtele M, Larsen NB, Ottosson DR, Storm P, Parmar M, et al (2022) Silk scaffolding drives self-assembly of functional and mature human brain organoids. Front Cell Dev Biol 10: 1023279

Tornavaca O, Chia M, Dufton N, Almagro LO, Conway DE, Randi AM, Schwartz MA, Matter K & Balda MS (2015) ZO-1 controls endothelial adherens junctions, cell-cell tension, angiogenesis, and barrier formation. Journal of Cell Biology 208: 821–838

Vabre P, Gatimel N, Moreau J, Gayrard V, Picard-Hagen N, Parinaud J & Leandri RD (2017) Environmental pollutants, a possible etiology for premature ovarian insufficiency: A narrative review of animal and human data. Environ Health 16: 37

Wagner M, Yoshihara M, Douagi I, Damdimopoulos A, Panula S, Petropoulos S, Lu H, Pettersson K, Palm K, Katayama S, et al (2020) Single-cell analysis of human ovarian cortex identifies distinct cell populations but no oogonial stem cells. Nat Commun 11: 1147

Wu T, Hu E, Xu S, Chen M, Guo P, Dai Z, Feng T, Zhou L, Tang W, Zhan L, et al (2021) clusterProfiler 4.0: A universal enrichment tool for interpreting omics data. Innovation 2: 100141

Yan J, Wu T, Zhang J, Gao Y, Wu JM & Wang S (2023) Revolutionizing the female reproductive system research using microfluidic chip platform. J Nanobiotechnology 21: 490

Yang J, Huang S, Cheng S, Jin Y, Zhang N & Wang Y (2021) Application of Ovarian Cancer Organoids in Precision Medicine: Key Challenges and Current Opportunities. Front Cell Dev Biol 9: 701429

Yoshino T, Suzuki T, Nagamatsu G, Yabukami H, Ikegaya M, Kishima M, Kita H, Imamura T, Nakashima K, Nishinakamura R, et al (2021) Generation of ovarian follicles from mouse pluripotent stem cells. Science (1979) 373: 298

Yu G, Wang LG, Han Y & He QY (2012) ClusterProfiler: An R package for comparing biological themes among gene clusters. OMICS 16: 284–287

Yu G, Wang LG, Yan GR & He QY (2015) DOSE: An R/Bioconductor package for disease ontology semantic and enrichment analysis. Bioinformatics 31: 608–609

Zhu A, Ibrahim JG & Love MI (2019) Heavy-Tailed prior distributions for sequence count data: Removing the noise and preserving large differences. Bioinformatics 35: 2084–2092

